# Targeting retrovirus-derived *Rtl8a* and *8b* causes late onset obesity and neurodevelopmental defects

**DOI:** 10.1101/2023.05.28.542606

**Authors:** Yoshifumi Fujioka, Hirosuke Shiura, Masayuki Ishii, Ryuichi Ono, Tsutomu Endo, Hiroshi Kiyonari, Yoshikazu Hirate, Hikaru Ito, Masami Kanai-Azuma, Takashi Kohda, Tomoko Kaneko-Ishino, Fumitoshi Ishino

## Abstract

Retrotransposon Gag-like (*RTL*) 8A, 8B and 8C are triplet genes of uncertain function that form a cluster on the X chromosome. They are eutherian-specific genes presumably derived from a certain retrovirus. Here, we demonstrate that *Rtl8a* and *Rtl8b* play an important role in growth and behavior via brain functions in the hypothalamus and prefrontal cortex. *Rtl8a* and *Rtl8b* double knockout (DKO) mice exhibited overgrowth due to hyperphagia from young adulthood and reduced social responses, increased apathy-like behavior. RTL8A and RTL8B proteins are localized to both the nucleus and cytoplasm of neurons presumably due to an N-terminal nuclear localization signal-like sequence. An increment in nucleus size was also detected in the neurons in the prefrontal cortex, suggesting neuronal dysfunction. These data give another strong evidence that retrovirus-derived acquired genes contributed to the establishment of the current eutherian developmental system in a wide variety of ways.

**Summary statement:** *Rtl8a* and *Rtl8b* double knockout mice exhibited late onset obesity and neurodevelopmental defects, demonstrating that these eutherian specific retrovirus-derived acquired genes encoding proteins with only 113 amino acids play important roles in the brain presumably via their functions in the hypothalamus and prefrontal cortex.

## Introductions

<u>R</u>etrotransposon Gag-like (*RTL,* also known as <u>s</u>ushi-ichi-<u>r</u>elated retrotransposon <u>h</u>omolog (*SIRH*)) genes encode proteins homologous to the sushi-ichi GAG (and sometimes also to POL) protein(s) (Ono *et al*., 2001, 2006; Charlier *et al*., 2001; Brandt *et al*., 2005; Youngson *et al*., 2005) and are presumed to be derived from a retrovirus (Imakawa *et al*., 2022). They play essential and/or important roles in eutherian development (Kaneko-Ishino and Ishino, 2012, 2015), such as paternally expressed 10 (*Peg10*), *Rtl1* (aka *Peg11*) and *Ldoc1* (aka *Rtl7* or *Sirh7*) in the placenta (Ono *et al*., 2006; Sekita *et al*., 2008; Naruse *et al*., 2014), *Rtl4* (aka *Sirh11* or *Zcchc16*) (Lim *et al*., 2013; Irie *et al*., 2015) and *Rtl1* (Kitazawa *et al*., 2022; Chou *et al*., 2022) in the brain and *Rtl6* (aka *Sirh3*) and *Rtl5* (aka *Sirh8*) in the brain innate immune system against bacteria and viruses as microglial genes (Irie *et al*., 2022). *Rtl1* is also important for muscle development (Kitazawa *et al*., 2021).

Biological roles of *RTL8A, 8B* and *8C* (aka *SIRH5, 6* and *4*) remain unknown although their involvement is recently indicated in two human disorders, Angelman syndrome (AS) and amyotrophic lateral sclerosis (ALS) because RTL8 as well as PEG10 proteins, another *RTL* gene product, increased in differentiated neurons from iPS of AS patients (Pandya *et al*., 2021) while RTL8 decreased but PEG10 increased in the differentiated neurons from iPS of ALS patients (Whiteley *et al*., 2021). In this work, we demonstrated that *Rtl8a* and *Rtl8b* double knockout (DKO) mice exhibit late onset obesity and apathy-like behavior but not abnormal physical functioning, implying the involvement of these genes in some human neurodevelopmental disease(s) but probably not ALS.

## Results and Discussion

### Conservation of *RTL8A-C* in eutherians

*RTL8A-C* (aka *SIRH5, 6* and *4*) forms a multi-gene cluster on the X chromosome. Its number (2-4, mostly 3, excluding pseudogenes) and amino acid (aa) sequence are well conserved (Table S1, red: dN/dS <0.1, pink: 0.1 ≤ dN/dS < 0.2), suggesting its important roles in eutherians. In most cases, *RTL8A-C* genes within the same species exhibit the highest homology to each other, suggesting independent gene conversion events in each species, therefore, they are not in a precise orthologous relationship among eutherians (Fig. S1).

In mice, *Rtl8a* and *Rtl8b* encode identical proteins with 113 aa, while *Rtl8c* encodes a protein with 112 aa exhibiting 70% aa homology to the former (Fig. 1A). They exhibit homology to suchi-ichi retrotransposon GAG (48.7 and 47.4% similarity and 34.2 and 28.9% identity, respectively). Mouse RTL8A-C (mRTL8A-C) exhibits a high aa sequence homology with those in other eutherian species on the whole (Table S1 and Fig. S2), but mRTL8A (also mRTL8B) has a rodent-specific nuclear localization signal (NLS)-like sequence (K-K/R-X-K/R) in the N-terminus (Fig. 1B) while mRTL8C has a unique leucine zipper motif (76-97 aa) (Figs. 1A, B and Fig. S2).

**Figure 1.**
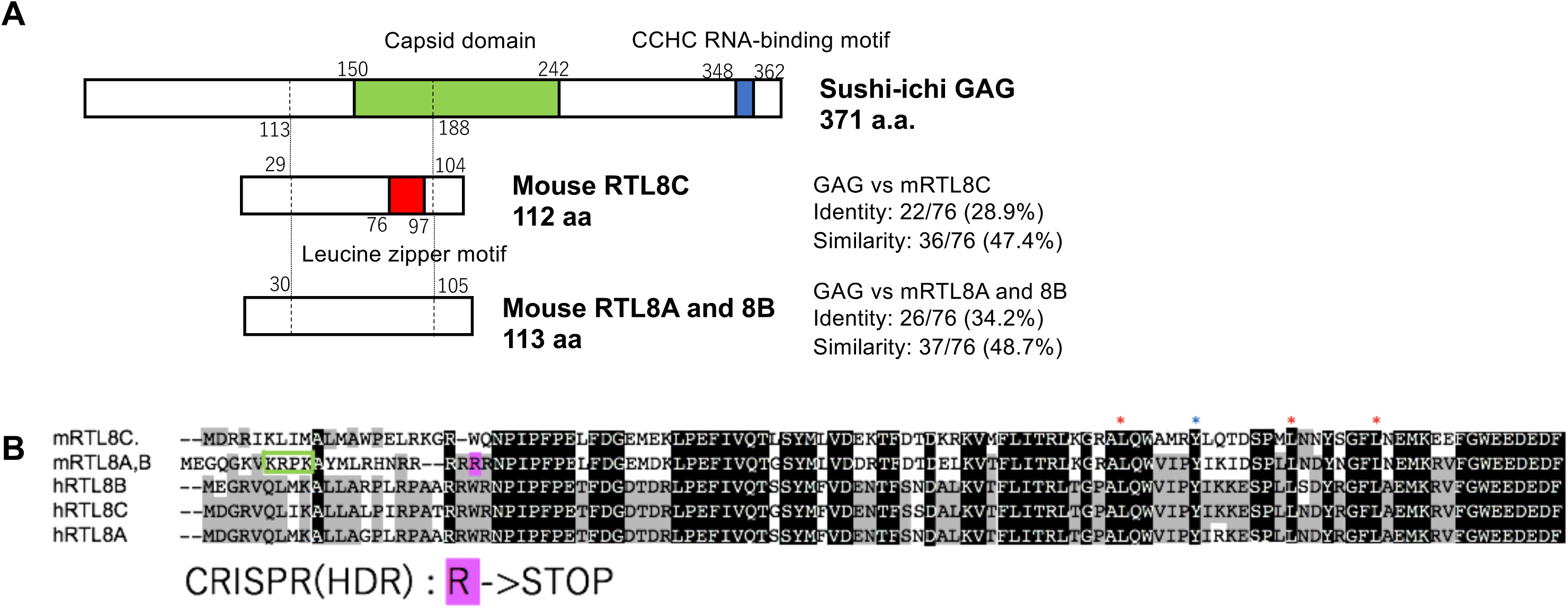
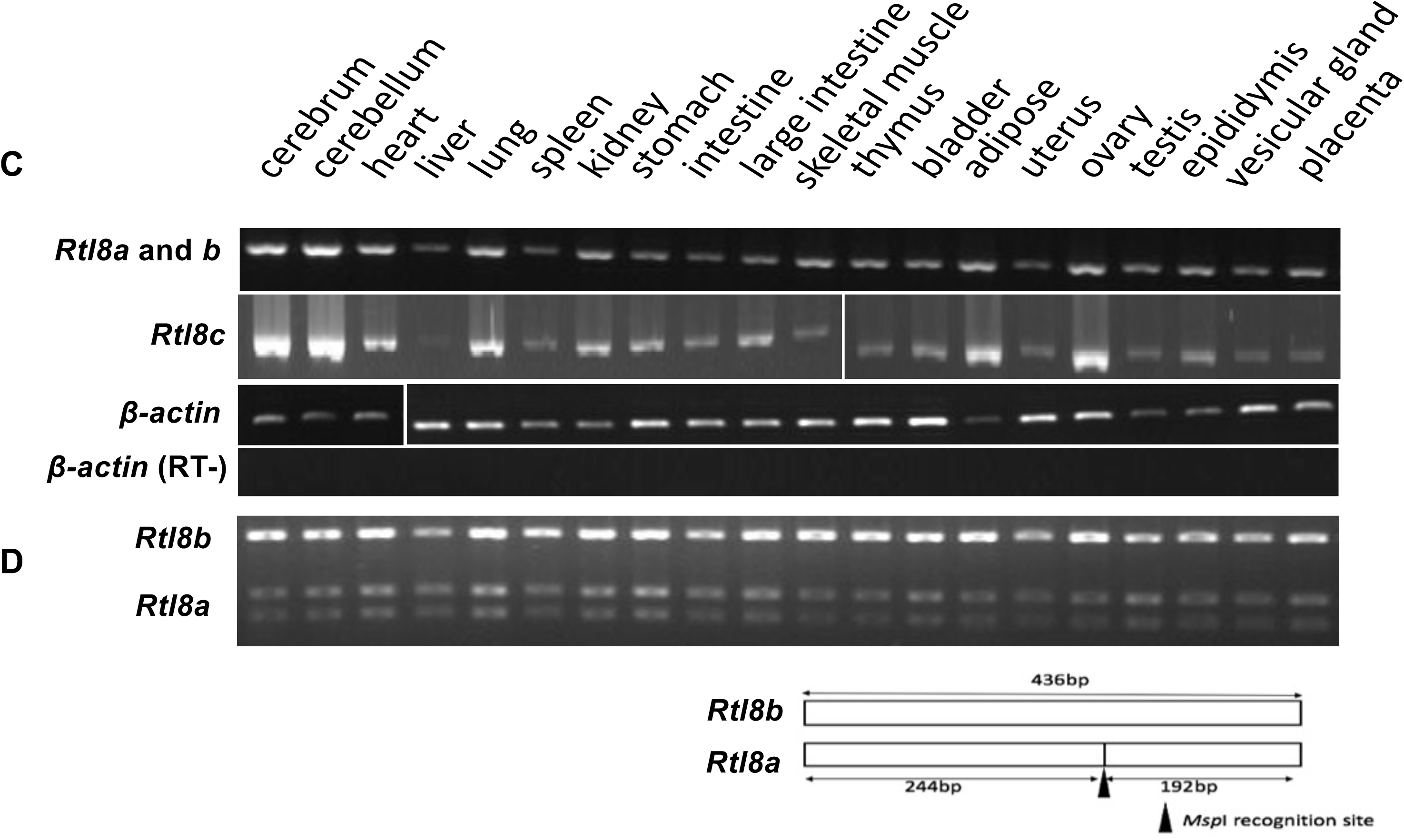
Characteristics of mouse RTL8A-C proteins. **(A)** Both mRTL8A and 8B comprise 113 aa with an identical aa sequence while mRTL8C comprises 112 aa including a unique leucine zipper motif (the red box) (see also Fig. S2). **(B**) Alignment of mouse and human RTL8A-C. The leucine zipper motif of mRTL8C is indicated by 4 asterisks (top). It should be noted that the second leucine (the purple asterisk) is unique to mRTL8C (see also Fig. S2). mRTL8A/8B has a rodent-specific nuclear localization signal (NLS)-like peptide in the N-terminus (the green box). A stop codon shown in pink was replaced with arginine (R) at 23 aa from the N-terminus in *Rtl8a* and *8b* DKO. **(C)** Expression profiles of *Rtl8a/8b* and *Rtl8c*. RT-PCR results in adult tissues and organs are shown. **(D)** RFLP analysis of *Rtl8a* and *8b*. The upper bands represent *Rtl8a* expression while the bottom two bands represent *Rtl8b* expression. The black arrowhead indicates the location of the recognition site of *Msp*1.

*Rtl8a*-*c* mRNAs exhibit higher expression in the cerebrum and cerebellum but ubiquitously expressed in all other tissues and organs except for no *Rtl8c* expression in the liver (Fig. 1C). *Rtl8a* and *Rtl8b* exhibit similar expression pattens estimated from a DNA polymorphism investigation using an *Msp*I recognition site (Fig. 1D). To address their biological functions, we first generated *Rtl8c* flox mice (Fig. S3) and then both *Rtl8a* and *Rtl8b* were knocked out by introducing a stop codon at a site 23 aa from the N-terminus of each gene using CRISPR/Cas9 mediated genome editing (Figs. S4, S5 and S6).

### Reproductive and growth abnormalities in *Rtl8a*/*8b* DKO mice

*Rtl8a*/*Rtl8b* heterozygous (hetero) DKO female mice exhibited abnormal maternal care and both female and hemizygous DKO male mice (hereafter called DKO males) exhibited overgrowth from the young adult stage. No lethality was observed in the DKO mice in the fetal and postnatal stages. Unexpectedly, one half of the hetero DKO females had delivery problems: most of their pups died soon after birth regardless of genotype (Fig. S7). The pups themselves, including homo DKO females, were viable when raised by surrogate mothers, suggesting abnormal maternal behavior of the hetero DKO mothers. The hetero DKO females may exhibit either normal or abnormal maternal behavior due to random X chromosome inactivation (XCI) in female mice (Fig. S7).

Both the hetero DKO females and DKO males gradually became overweight after 8 w associated with the increased food intake and this trend continued subsequently (Fig. 2A and Fig. S8), suggesting certain dysfunction of the hypothalamus. One of the maximal weights of the hetero DKO females reached 75 g at 80 w compared to the 34 g of a WT littermate female (Fig. 2B).

**Figure 2.**
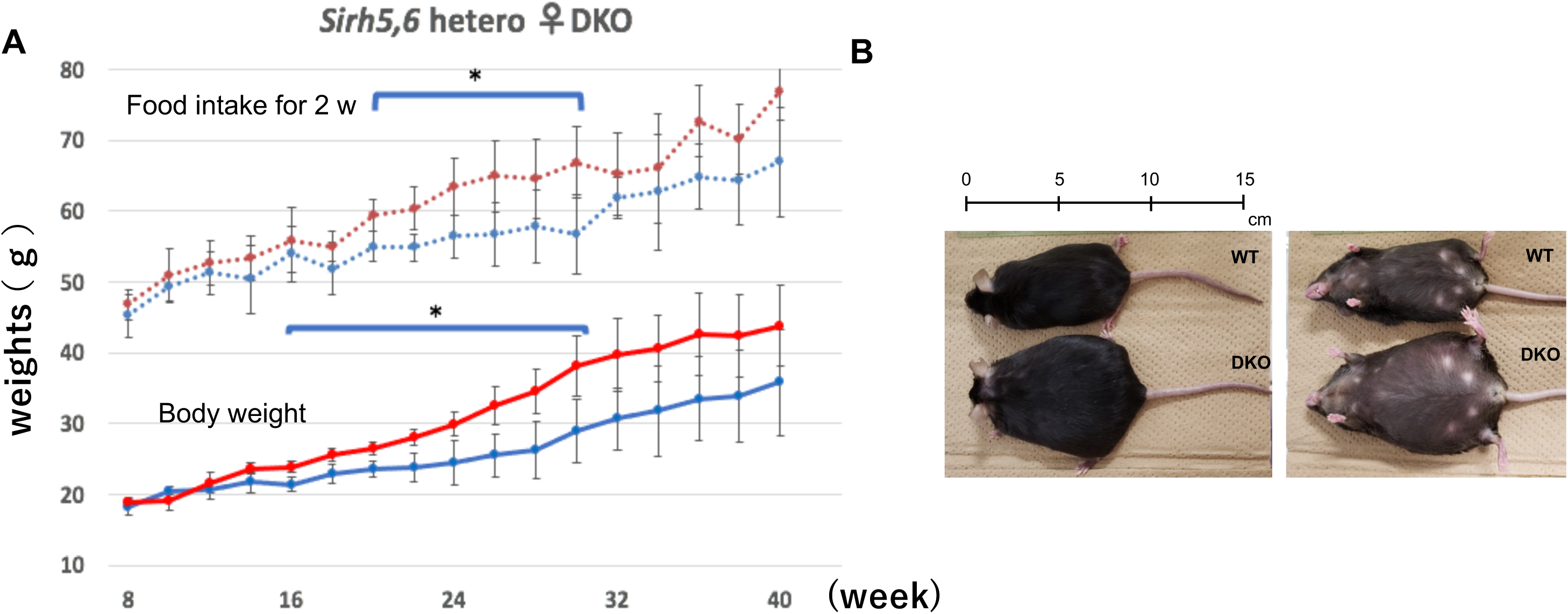
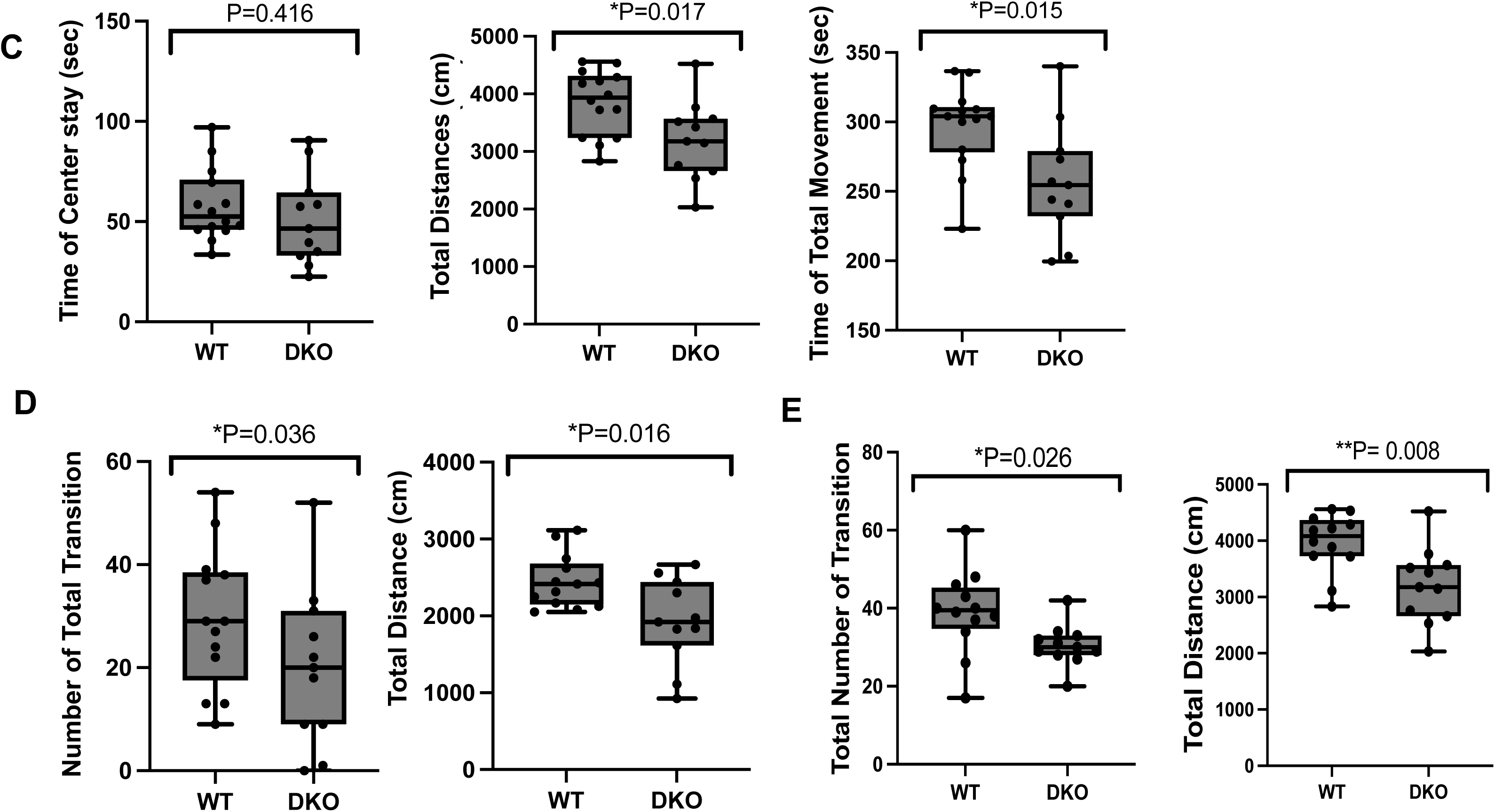
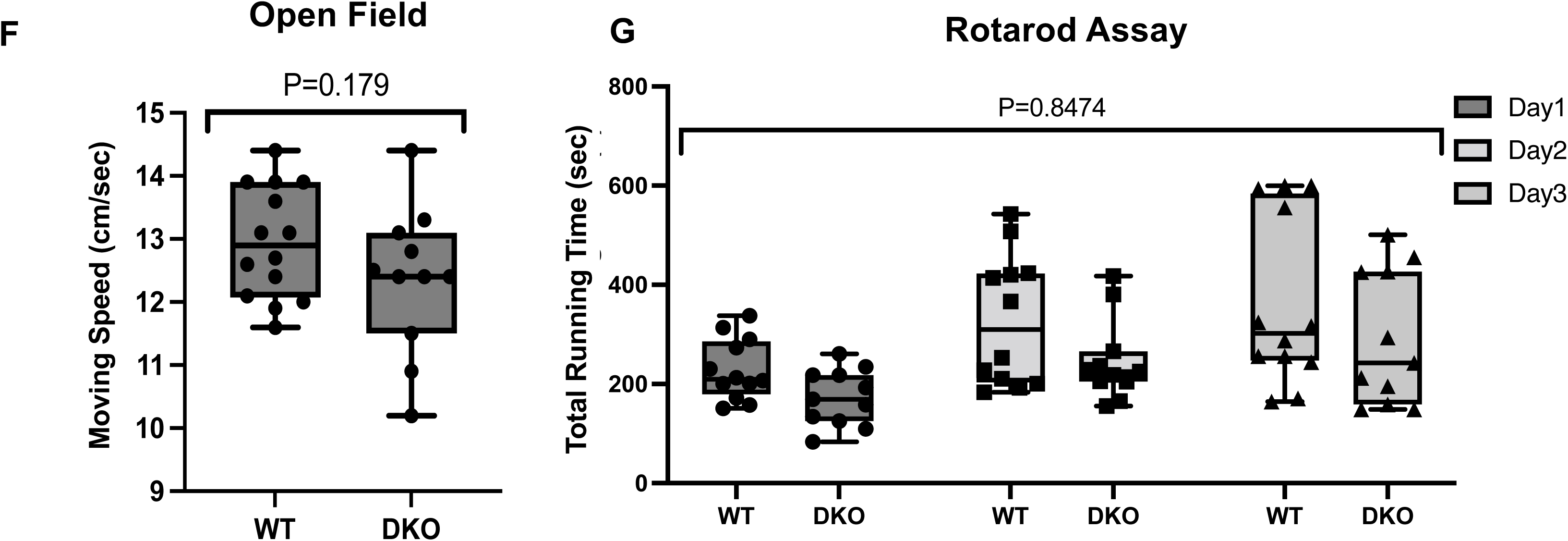
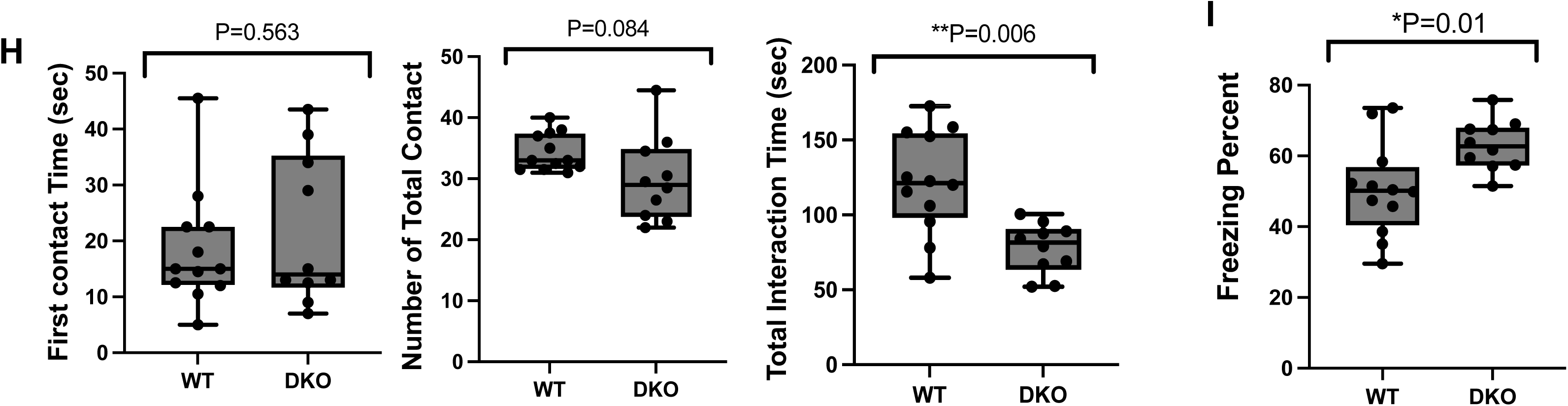
Obesity and abnormal behaviors of *Rtl8a* and *8b* DKO. **(A)** Body weight (bottom) and food intake every two weeks (top) of the WT (n = 5) and hetero DKO female (n =5) mice are shown. Blue: WT, Red: DKO. * P < 0.05. **(B)** One example of the maximal weight of *Rtl8a* and *8b* hetero DKO females (bottom, 75 g) compared with a WT littermate female (top, 34 g) at 80 weeks. **(C-I)** Behavior tests. Reduced locomotive activity was observed in the OF test **(C)** (WT: n = 12, DKO: n = 10, t-test), L/D transmission test **(D)** (WT: n = 14, DKO: n = 11, t-test) and EPM test **(E)** (WT: n = 12, DKO: n = 11, t-test). *P < 0.05, **P < 0.01. Normal physical capability was observed in the OF test **(F)** (WT: n=14, DKO: n=11, t-test) and rotarod assay **(G)** (WT: n = 12, DKO: n = 11, two-way ANOVA 0.8474). Reduced social activity was observed in the social behavior test **(H)** (WT: n = 12, DKO: n = 10, t-test). Increased apathy-like behavior was observed in the tail suspension test **(I)** (WT: n = 12, DKO: n = 10, t-test). *P < 0.05, **P < 0.01.

### Abnormal behaviors in *Rtl8a/8b* DKO mice

The DKO mice exhibited reduced locomotive activity, decreased social responses and increased apathy. In the open field (OF: Fig. 2C), light/dark transition (L/D: Fig. 2D, left) and elevated plus-maze tests (EPM: Fig. 2E, right), the DKO males exhibited reduced locomotive activity in terms of the total distance (3953 vs 3191 cm in OP; t-test 0.017, 2373 vs 1925 cm; t-test 0.016 in L/D, and 1961 vs 1653 cm; t-test 0.023 in EPM), total movement (296 vs 257 sec; t-test 0.015 in OF) and number of transitions (29.7 vs 17.9 times; t-test 0.036 in L/D and 37.5 vs 30.4 times; t-test 0.046 in EPM). The other elements of these tests were normal (Figs. S9 and S10) including normal moving speed (OF: Fig. 2F) and a normal score in the rotarod test (Fig. 2G), suggesting their normal physical functioning.

In the social activity test, they had similar first contact time and the number of total contacts compared with their WT littermates, but the total interaction time was reduced (121.5 vs 81.6 sec; t-test 0.006), suggesting a reduction in social activity (Fig. 2H). In the tail suspension test (Fig. 2I), their freezing time was increased (50.3 vs 63 sec; t-test 0.01), suggesting increased apathy. We did not observe any abnormality in the fear conditioning test except for lower locomotive activity (Fig. S11). We also found that some DKO males were severely injured by attacks by other DKO males, while certain other DKO males suffered severe self-inflicted injuries, biting and scratching themselves during normal breeding, suggesting that the loss of RTL8A/8B somehow affects the neuronal functions in the brain.

### mRTL8A/8B expression in the hypothalamus

mRTL8A/8B proteins were detected in certain specific regions, such as the hypothalamus, frontal/temporal cortex, striatum and olfactory bulb (Figs. 3A-E and Fig. S12). For detection of the mRTL8A/8B proteins, we used an in-house anti-mRTL8A/8B antibody by immunization with their unique N-terminal peptides (Fig. 1B and Fig. S2). We confirmed the antibody specificity by checking whether WT signals disappeared in the DKO brain (Figs. 3A and B).

**Figure 3.**
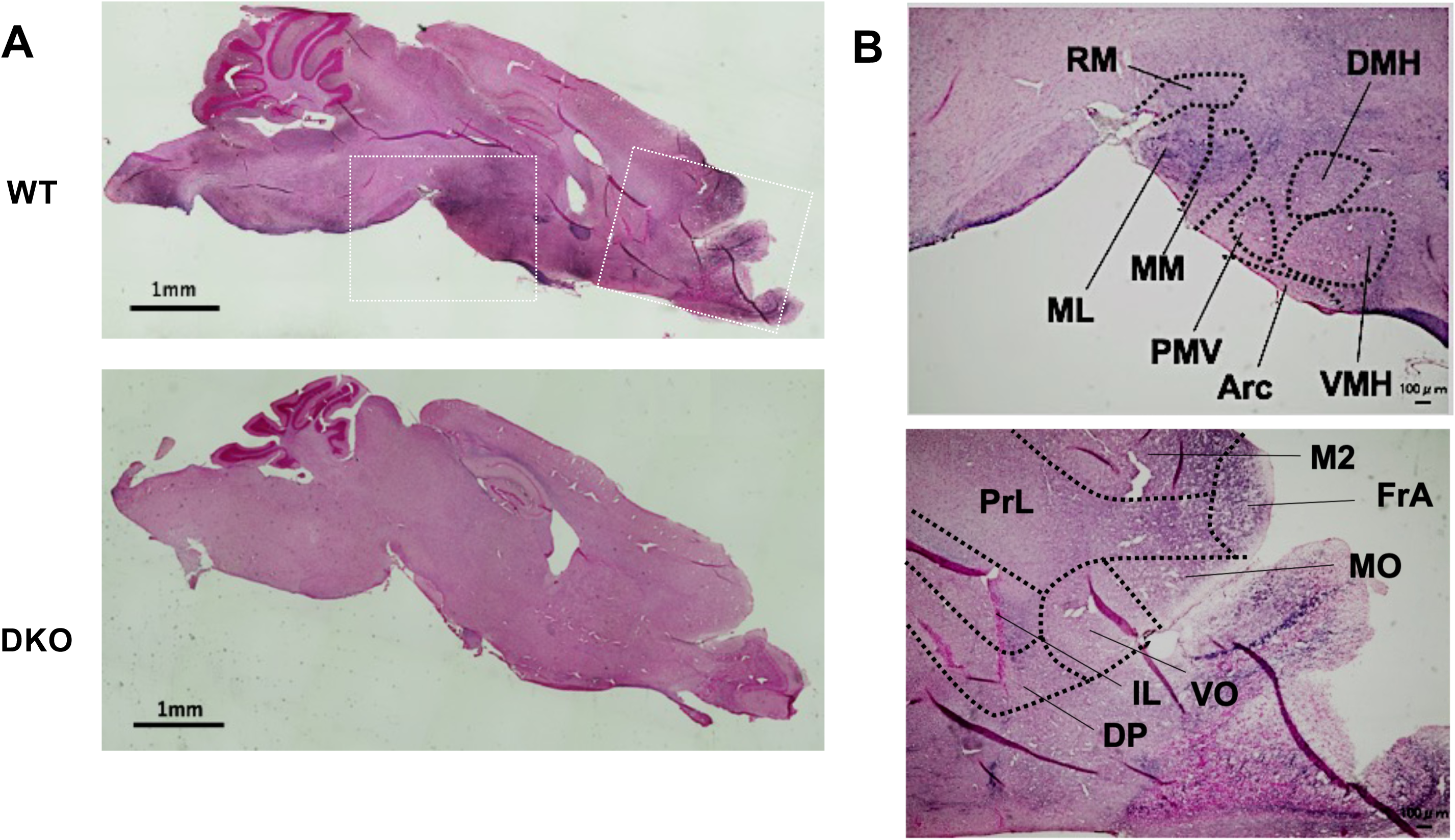
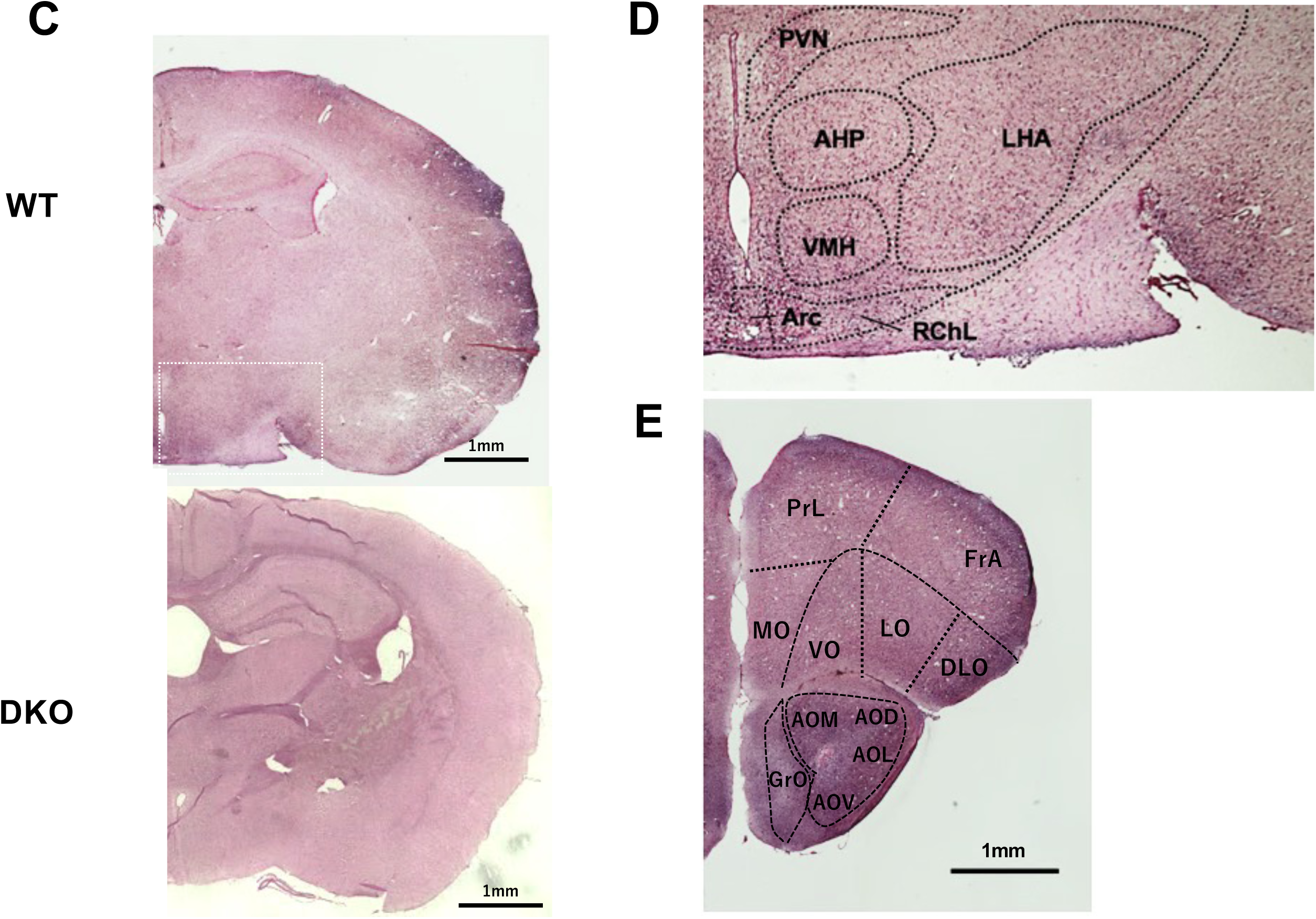
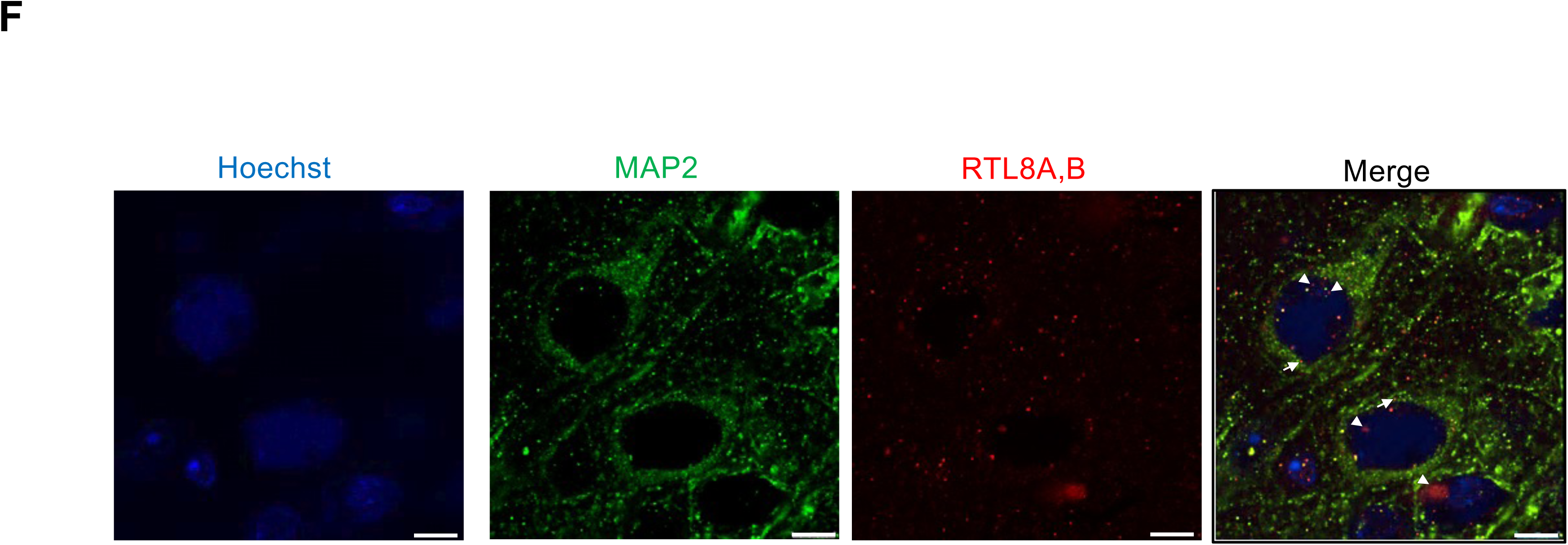
Immunostaining of mRTL8A and 8B proteins in the brain. **(A)** Sagittal section images of the WT (top) and DKO (bottom) mice (each at 20 w). mRTL8A/8B protein was detected in the frontal cortex and hypothalamus (n=5). **(B)** Enlarged areas surrounded by white squares in **(A).** (Top) Arc: Arcuate nucleus, DMH: Dorsomedial hypothalamic nuclei, ML: medial mammillary nucleus, MM: Medial mammillary nucleus, PMV: premammillary nucleus, ventral part, RM: retromammillary nucleus, VMH: Ventromedial hypothalamic nuclei. (Bottom) DP: dorsal peduncular cortex, FrA: frontal association, IL: infralimbic cortex, M2: secondary motor cortex, MO: medial orbital cortex, PrL: Prelimbic cortex, VO: ventral orbital cortex. **(C)** Coronal section images of the WT (top) and DKO (bottom) mice (each at 20 w). **(D)** An enlarged areas surrounded by white squares in **(C)**. AHP: anterior hypothalamus nucleus, LHA: lateral hypothalamic area. PVN: paraventricular nucleus, RChL: Retrochiasmatic area, lateral part. **(E)** A coronal section images of the PFC and olfactory bulb regions of WT mice (at 10 w). AOD: anterior olfactory area, dorsal part, AOL: anterior olfactory area lateral part, AOM: anterior olfactory area medial part, AOV: anterior olfactory area ventral part, DLO: dorsolateral orbital cortex, FrA: frontal association, GrO: granule cell layer of the olfactory bulb, LO: lateral orbital cortex, MO: medial orbital cortex, PrL: prelimbic cortex, VO: ventral orbital cortex. **(F)** Coimmunofluorescence staining images of MAP2 and RTL8A/8B in the FrA region in WT (sagittal sections, each at 20 w, n=2). White arrowheads and arrows indicate the RTL8A/8B signals in the nuclei and cytoplasm, respectively. Green: MAP2, Red: mRTL8A/8B blue: Hoechst33342. Scale bars: 8 μm.

mRTL8A/8B were detected in the most hypothalamic regions (Figs. 3A, B, E and F), including several important hypothalamic nuclei related to feeding behavior, such as the arcuate nucleus (Arc) (Lanfray and Richard, 2017), dorsomedial and ventromedial hypothalamic nuclei (DMH and VMH) (Merkestein *et al*., 2014), paraventricular nucleus (PVN) (Yousefvand and Hamidi, 2020), anterior hypothalamic area, posterior part (AHP) and lateral hypothalamus (LHA) (Figs. 3E and F). In addition, they were detected in the nucleus accumbens (Acb) in the ventral striatum (Fig. S12) and medial forebrain bundle integrated in the LHA that connects the Acb to the ventral tegmental area (VTA) in the midbrain, two important control centers for feeding behavior (Atasoy *et al*., 2012). In the DMH region, mRTL8A/8B exhibited an immunostaining image pattern similar to that obtained using Neurofilament (Fig. S12), suggesting their neuronal expression, consistent with a previous reports that human RTL8A-C proteins are expressed in the iNeurons differentiated from AS and ALS patients’ iPS cells (Pandya *et al*., 2021, Whiteley *et al*., 2021).

### mRTL8A/8B expression in the brain

mRTL8A/8B were also detected in a wide area of the prefrontal cortex (PFC). PFC includes the orbital cortex, prelimbic cortex (PrL) and frontal association (FrA) (Figs. 3C and D). The medial orbital cortex (MO) is a part of the orbitofrontal cortex (OFC) that plays an important role in the regulation of feeding behavior (Seabrook and Borgland, 2020). The OFC, a ventral subregion of the PFC (Szczepanski and Knight, 2014), is also involved in the integration of sensory information, emotion processing, decision-making and behavioral flexibility (Hare *et al*., 2009; Hare and Duman, 2020). The PrL is involved in depression-like despair behavior (Pizzagalli and Roberts, 2022) and maintaining attention to category-relevant information, and flexibly updates category representations (Broschard *et al*., 2021), while the FrA is engaged in stimulus integration during associative learning (Nakayama *et al*., 2015). The FrA receives projection from the insular (IC) and perirhinal (PRh) cortices in the temporal lobe. The high mRTL8A/8B expression was also detected in these regions (Fig. S13) that are activated by context exposure and shock, respectively (Nakayama *et al*., 2015).

Functional impairment of all these regions may explain the reduced locomotive activity and social interaction as well as the increased apathy-like behavior.

Co-immunostaining experiment of mRTL8A/8B with a neuronal marker, microtubule associated protein (MAP) 2, demonstrated that these proteins are located both in the nucleus and cytoplasm of neurons (Fig. 3F and Fig. S14). It is consistent with the fact that they have the N-terminal NLS-like signals (Fig. 1B) and also with the fact that human RTL8A/8B are involved in the nuclear transition of UBQLN2 that is necessary for nuclear protein quality control (Mohan *et al*., 2022). In addition, certain neuronal deformation was observed in the medial PFC (mPFC) (Figs. 3C, D and E).

Immunofluorescence staining with anti-MAP2 antibody also demonstrated that the nuclei of the layer 2/3 pyramidal cells in the FrA were significantly increased in size (Figs. 4A and B, Fig. S15) in the 20 w adult brain (Fig. 4B, Fig. S16). This was not observed in the other cortical regions without mRTL8A/8B expression (Fig. 3C), such as the primary somatosensory cortex, shoulder region (S1Sh) (Figs. 4C and D, Figs. S17 and S18). It has been reported that the pyramidal cells in the mPFC moderate stress induced depressive behaviors (Shrestha *et al*., 2015), suggesting that this neuronal change is related to the apathy-like behavior observed in the DKO mice possibly due to abnormal nuclear protein quality above described.

**Figure 4.**
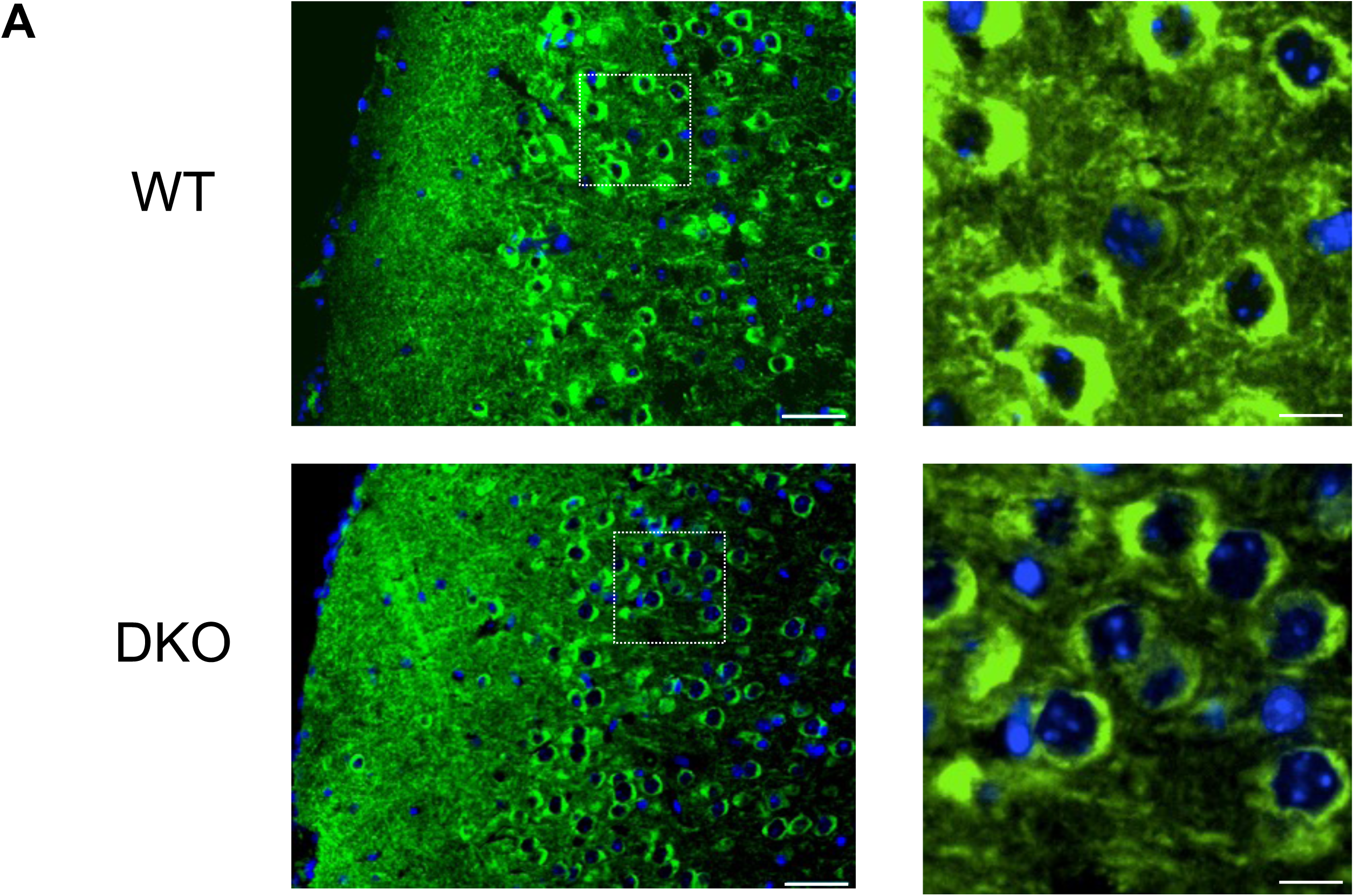
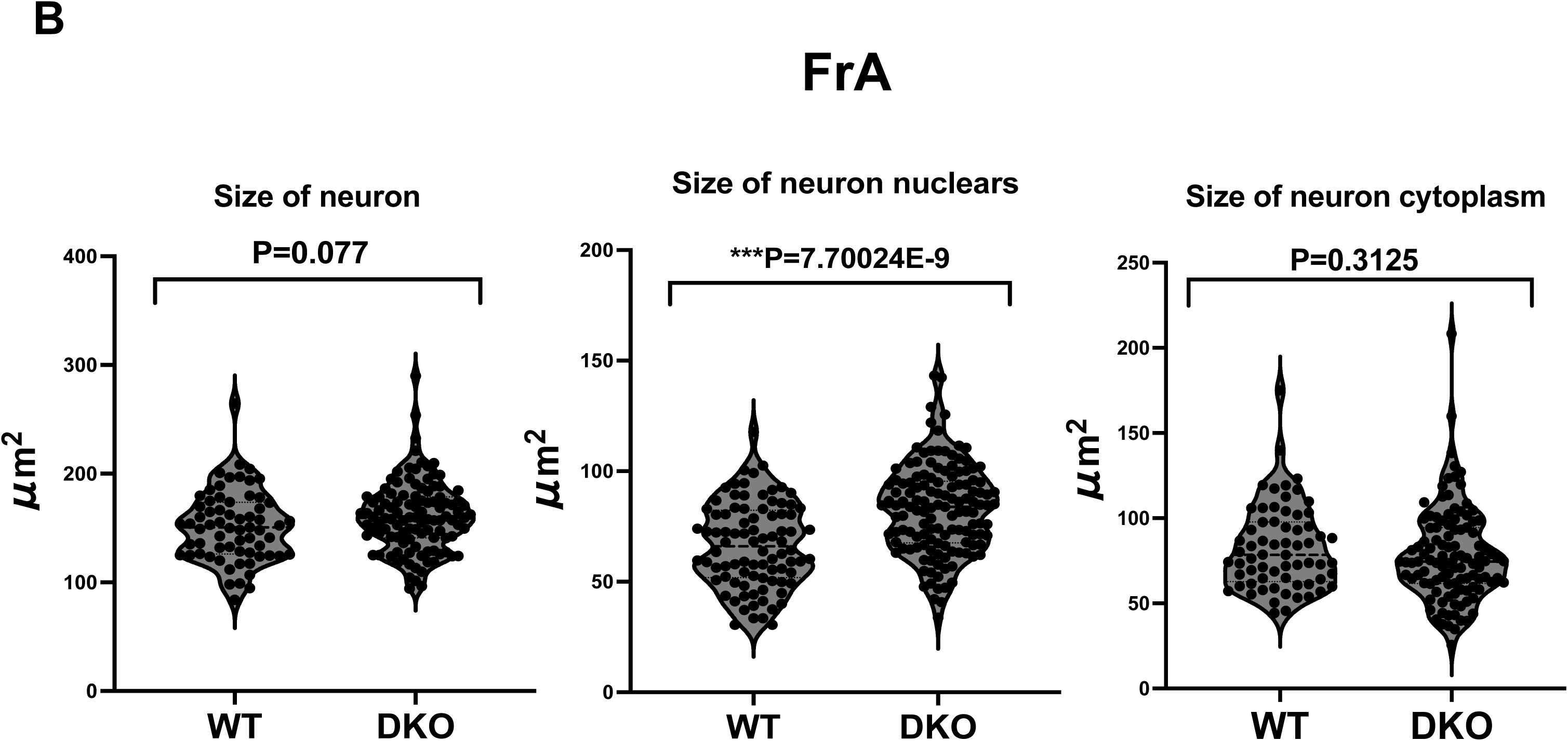
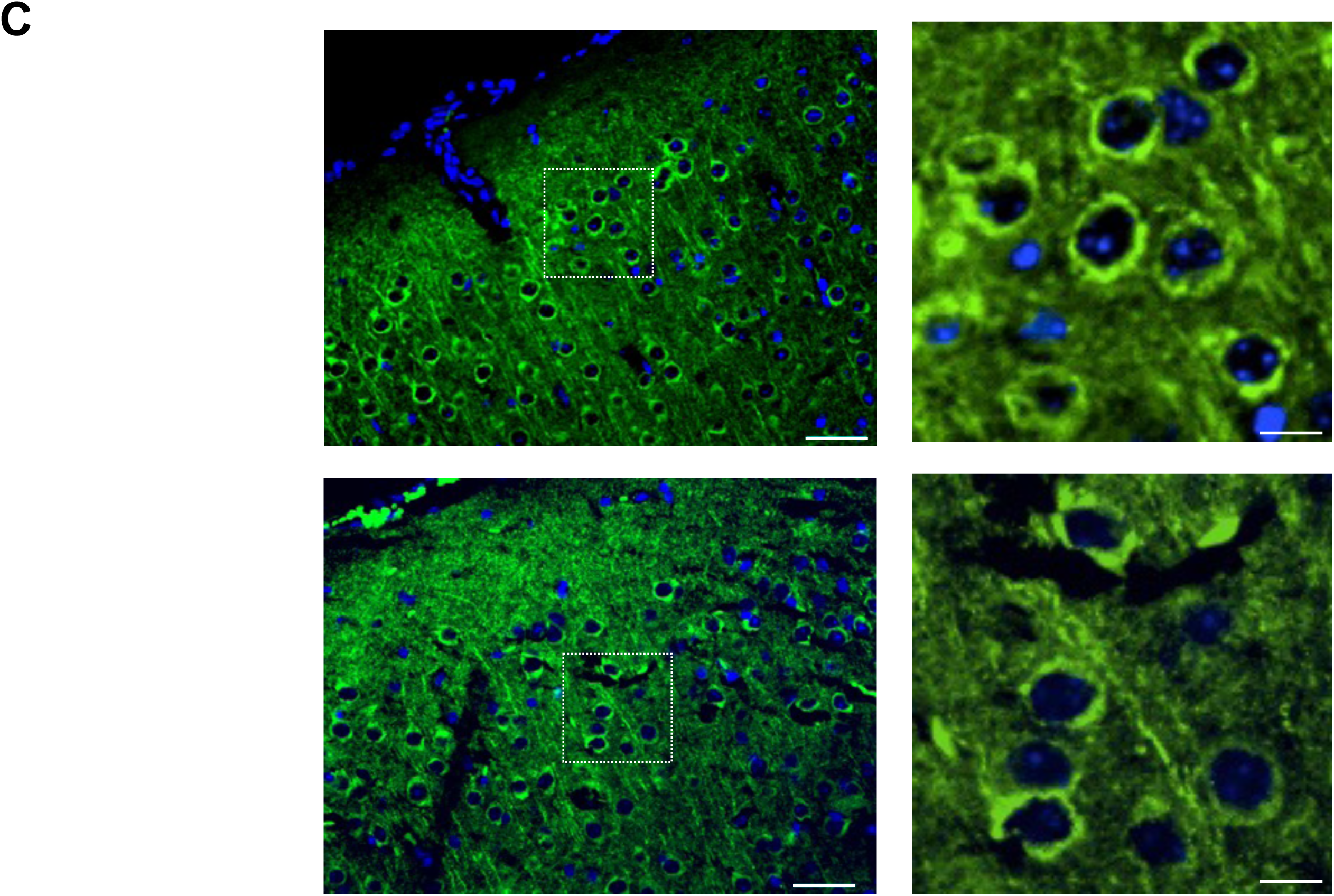
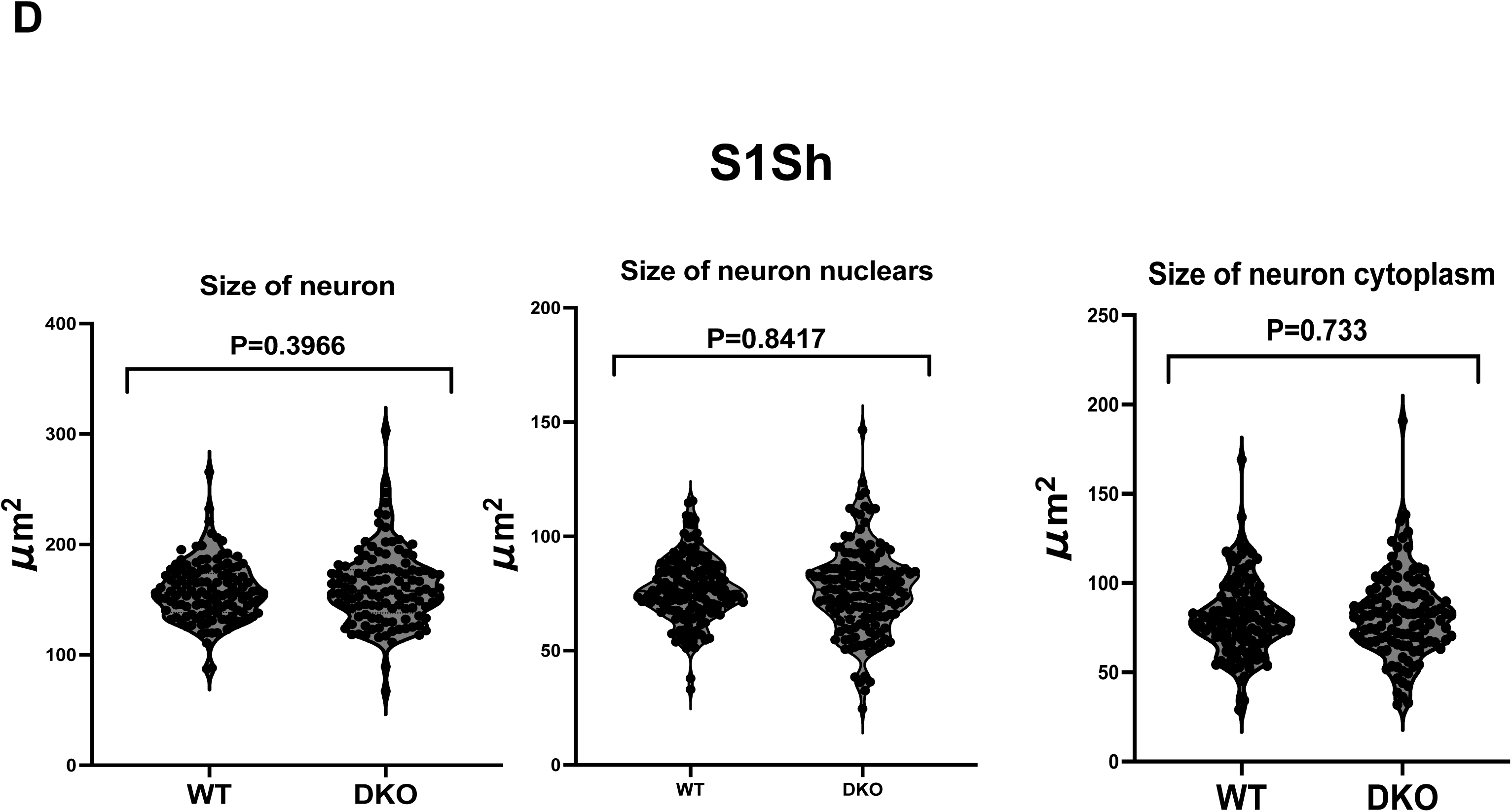
Immunofluorescence staining of FrA in WT and DKO mice. **(A)** Left: Immunofluorescence staining images of MAP2 in the FrA region in WT and DKO mice (sagittal sections, each at 20 w, n=2). Right: Enlarged images of the dashed box areas in the left pictures. Green: MAP2, blue: DAPI. Scale bars: 40 μm (left) and 10 μm (right), respectively. **(B)** The sizes of the neuronal cells and their nuclei in (A) were measured by a blind observer using ImageJ. Three individuals from each of the WT (blue) and KO (red) mice were examined for the measurement of a total of 65 and 109 neurons, 88 and 132 neuronal nuclei, and 65 and 109 neuronal cytoplasm samples, respectively. All measurements were statistically analyzed with Student’s t-test and the results are displayed as Violin Plots. *P < 0.05, **P < 0.01, ***P< 0.001. **(C)** Left: Immunofluorescence staining images of MAP2 in the S1Sh region in the WT and DKO mice (Sagittal sections, each at 20 w). Right: Enlarged images of the dashed box areas in the left pictures. Green: MAP2, red: DAPI. Scale bars: 40 μm (left) and 10 μm (right), respectively. **(D)** The sizes of the neuronal cells and their nuclei in (C) were measured by a blind observer using ImageJ. Three individuals from each of the WT (blue) and KO (red) mice were examined for the measurement of a total 125 and 116 neurons, 163 and 149 neuronal nuclei and 125 and 116 neuronal cytoplasm samples, respectively. All measurements were statistically analyzed with Student’s t-test and the results are displayed as Violin Plots. *P < 0.05, **P < 0.01, ***P< 0.001.

### Biological significance of *RTL8A*/*8B* and possible link to human disease(s)

This study demonstrates that retrovirus-derived *Rtl8a*/8*b* play important roles in the brain via regulating certain basic behaviors, such as food intake, maternal care and emotions (Figs. 2 and 3, and Figs. S7 and S8) with their 113 aa proteins (Fig. 1A). This explains why *RTL8A-C* is well conserved in eutherians (Table S1).

Interestingly, the overgrowth due to an increment of food intake after 8w (Figs. 2A and B) as well as reduced social interaction (Fig. 2H) and increased apathy (Fig. 2I) in DKO are similar to those observed in the late Prader-Willi syndrome (PWS) patients that present late onset obesity and ASD-like symptoms from the juvenile period (Nicholls and Knepper, 2001). PWS is a neurodevelopmental genomic imprinting disease caused by maternal duplication of human chromosome 15q11-q13. Interestingly, the involvement of *RTL8* is suggested in AS (Pandya *et al*., 2021) that is caused by paternal duplication of the same imprinted region (Nicholls and Knepper, 2001). Given that human RTL8A-C proteins are direct targets of UBE3A (Pandya *et al*., 2021), the RTL8A-C level should be reduced in the PWS patients because of double dosage of maternally expressed *UBE3A* (Nicholls and Knepper, 2001). Therefore, it is of interest to elucidate the involvement of RTL8 in PWS in future. However, the DKO mice had a normal score in the rotarod test (Fig 2G), suggesting that downregulation of these genes is not correlated with the phenotype of ALS (Whiteley *et al*., 2021).

In addition to *RTL8A-C,* at least *RTL1* and *RTL4* are involved in neurodevelopmental disorders, such as KOS14/TS14 (Sekita *et al*., 2008; Kagami *et al*., 2008, 2015; Ioannides *et al*., 2014; Kitazawa *et al*., 2021; Chou *et al.,* 2022) and ASD (Lim *et al*., 2013; Irie *et al*., 2015). Therefore, eutherian-specific retroviral *Gag*-derived genes should have garnered more attention as important targets in human neurodevelopmental diseases.

## Materials and Methods

### Animals

All of the animal experiments were reviewed and approved by the Institutional Animal Care and Use Committee of RIKEN Kobe Branch, Tokai University and Tokyo Medical and Dental University (TMDU) and were performed in accordance with the RIKEN Guiding Principles for the Care and Use of Laboratory Animals, and the Guideline for the Care and Use of Laboratory Animals of Tokai University and TMDU.

### Estimation of the pairwise dN/dS ratio

Conversion of the protein sequence alignment created with the MAFFT program (https://mafft.cbrc.jp/alignment/server/index.html) into the corresponding codon alignment was performed with the PAL2NAL program (www.bork.embl.de/pal2nal/) (Suyama *et al*, 2006). The PAL2NAL program simultaneously calculated the nonsynonymous/synonymous substitution rate ratio (dN/dS) by the CodeML program (runmode: −2) in PAML (Xu & Yang, 2013). An aa sequence phylogenic tree was constructed with MEGA5 using the Maximum Likelihood method based on the JTT matrix based model (Tamura *et al*. 2011). The bootstrap consensus tree inferred from 1000 replicates is taken to represent the evolutionary history of the taxa analyzed. Branches corresponding to partitions reproduced in less than 50% bootstrap replicates are collapsed. The percentage of replicate trees in which the associated taxa were clustered together in the bootstrap test (1000 replicates) are shown next to the branches. Initial tree(s) for the heuristic search were obtained automatically as follows. When the number of common sites was < 100 or less than one fourth of the total number of sites, the maximum parsimony method was used; otherwise the BIONJ method with MCL distance matrix was used. The tree is drawn to scale, with branch lengths measured in terms of the number of substitutions per site. The analysis involved 10 aa sequences. All positions containing gaps and missing data were eliminated. There was a total of 104 positions in the final dataset.

The *RTL8A, 8B* and *8C* genome sequences used for the dN/dS analysis (Table S1) were as follows: Human: NC_000023.11[c135052108-135051767, c135022459-135022118, 135032384-135032725]; Chimpanzee: NC_036902.1[c130245552-130245211, c130216735-130216394, 130226101-130226442]; Marmoset: NC_013918.1[122938486-122938827, c122959788-122959447, c123007085-123006744]; Mouse: NC_000086.8[c52645588-52645247, c52672265-52671924, 52610013-52610351]; Cow: NC_037357.1[18783020-18783361, 18794515-18794856, 18824715-18825056]; Rabbit: NC_013690.1[109509895-109510236, 109553752-109554093, 109580009-10958035]; Dog: NC_049780.1 [106817683-106818024, 106834142-106834483, 106843724-106844065, 106859125-106859466]; Elephant: NW_003573520.1[4927338-4927679, c 4938179-4937838] and NW_003575145.1[c434-87].

### RT-PCR

RT-PCR was performed using cDNA. cDNA was synthesized from 1 μg of total RNA using SuperScript Ⅲ Reverse Transcriptase (Invitrogen). RNA was extracted from adult tissues at 19 weeks, including the cerebrum, cerebellum, heart, liver, lung, spleen, kidney, stomach, intestine, large intestine, skeletal muscle, thymus, bladder, adipose, uterus, ovary, testis, epididymis and vesicular gland as well as day 9.5 placenta by treatment with TRIzol Reagent (Life Technologies). Ten ng of cDNA were mixed with 1 x ExTaq buffer (TAKARA), 2.5 mM of each dNTP 2 μl, primer 0.2 μl (200 pmol/μl) and ExTaq HS 0.1 μl (TAKARA), and PCR analysis was performed under the following conditions. *Rtl8a, b* and *c*: 95℃ 1min, 32 (*Rtl8a* and *b*) and 31 (*Rtl8c*) cycles at 96℃ for 10 s, 72℃ for 30 s and 72℃ for 90 s. *β-actin*: 95℃ 1min, 26 cycles at 96℃ for 10 s, 72℃ for 30 s, and 72℃ for 90 s using a C1000 Touch thermal cycler (Biorad). The following primer sets were used. Rtl8a, b F1: tcccactgttgacagctcag and Rtl8a, b R1: ggggctactgttggaaaa, and Rtl8c F1: tatgcgctatctaaagacagac and Rtl8c R1: ggcttctgtacaggtagaggcagac. The RFLP (restriction fragment length polymorphism) experiment was carried out using *Msp*I, and the results were confirmed by electrophoresis.

### Generation of *Rtl8c* flox mice

A targeting construct spanning the two exons of *Rtl8c* was used to generate *Rtl8c* flox mice (Accession No. CDB0788K: https://large.riken.jp/distribution/mutant-list.html). Briefly, the construct included two loxP sites upstream of exon 1 and downstream of exon 2, and a frt-neo-frt cassette downstream of the latter loxP site. After linearizing, the construct was electroporated into TT2 ES cells (C57BL/6 × CBA genetic background) (Yagi *et al*, 1993). Two ES clones were then screened by PCR and Southern blotting. One 5′ and one 3′ probe, both of which were outside of targeting construct, were used in Southern blotting. The correct targeting clones were then used to generate chimera founder lines. Germline transmission was observed in two chimeric mice.

The *Rtl8c* flox alleles were checked by electrophoresis of the PCR amplification using the following primers. WT forward: GGCGACAACCAAGGTTTTTA and reverse: GGGTCTGCTTCTCTTTGCTG, flox allele forward: AAATAGGCGTATCACGAGGC and reverse: GGGTCTGCTTCTCTTTGCTG. The PCR cycle was 96℃ 1min, 35 cycles at 96℃ for 15 s, 60℃ for 30 s and 72℃ for 30 s. and 72℃ 2min.

### Generation of *Rtl8a/Rtl8b* DKO mice

The DKO mice were generated by referring to our previous report (Ono *et al,* 2015) as follows. The plasmids expressing *hCas9* and sgRNA were prepared by ligating oligos into the *Bbs*I site of pX330 (Addgene plasmid #42230). The 20 bp sgRNA recognition sequences was: *Rtl8a, b*-sgRNA (5′-ACGGGATGGGGTTCCGCCGA-3′). The sgRNA targets both *Rtl8a* and *Rtl8b*.

To produce the *hCas9* mRNA, the T7 promoter was added to the *hCas9* coding region of the pX330 plasmid by PCR amplification, as previously reported (Wang *et al*, 2013). The T7-*Cas9* PCR product was gel purified and employed as the template for *in vitro* transcription (IVT) using a mMESSAGE mMACHINE T7 ULTRA kit (Life Technologies). The T7 promoter was added to the *Rtl8a, b*-sgRNA region of the pX330 plasmid by PCR amplification using the following primers. *Rtl8a, b*-sgRNA-F (5′-TGTAATACGACTCACTATAGGGACGGGATGGGGTTCCGCCGA-3′), *Rtl8a, b*-sgRNA-R (5′-AAAAGCACCGACTCGGTGCC-3′). The T7-sgRNA PCR product was gel-purified and employed as the template for IVT using a MEGAshortscript T7 kit (Life Technologies). Both the *hCas9* mRNA and *Rtl8a, b*-sgRNA were DNase-treated to eliminate template DNA, purified using a MEGAclear kit (Life Technologies), and eluted into RNase-free water.

C57BL/6N female mice were superovulated and *in vitro fertilization* was carried out using *Rtl8c* flox mouse sperm. The synthesized *hCas9* mRNA (50 ng/μl) and *Rtl8a, b*-sgRNA (25 ng/μl) with oligo DNA (50 ng/μl) were injected into the cytoplasm of fertilized eggs at the indicated concentration. The eggs were cultivated in KSOM overnight, then transferred into the oviducts of pseudopregnant ICR females. The oligo DNA designed as the HDR (homology-directed repair) donor induced a nonsense mutation in both *Rtl8a* and *Rtl8b* and harbored an accompanying *Afl*II recognition site used for PCR-RFLP (Restriction Fragment Length Polymorphism) genotyping. The sequence of the oligo DNA was: 5′-AAGGCCAAGGCAAGGTAAAGAGGCCGAAGGCCTACATGCTCAGGCACAACAGGCGGCGCCCTTAAGGGAACCCCATCCCGTTTCCAGAGCTGTTTGATGGCGAGATGGACAAGCTCCCGGAGTTCA-3′.

### Genotyping of *Rtl8a* and *8b* DKO mice

The genotype was determined by PCR-RFLP analysis. The following primers were used for PCR amplifications, *Rtl8a*: 5′-GGACTGGCGCCTGAAATAGC-3′ and 5′-GCACAATCACCACCTCTTGAACA-3′, *Rtl8b*: 5′-CCACCCCTTAAACATTCTCCTGG-3′ and 5′-AGATCGAACATCAGGCCATGAAC-3′. The PCR products were digested with *Afl*II and subjected to agarose gel electrophoresis.

### Behavioral Analysis

Behavioral analysis was performed as previously described (Irie *et al*, 2014; Kitazawa *et al,* 2021). Briefly, both DKO and littermate WT male mice (8-20 w) were analyzed in the following order, Light and Dark transition, Open field, Social Interaction, Elevated plus maze, Rotarod test, Tail suspension and Fear condition tests.

#### Light and Dark transition test

The apparatus used for the light/dark transition test comprised a cage (21 × 42 × 25 cm) divided into two sections of equal size by a partition with a door (O’hara & Co., LTD.). One chamber was brightly illuminated (400 lux), whereas the other chamber was dark. Each mouse was placed into the dark side and allowed to move freely between the two chambers with the door open for 10 min. The total number of transitions between the chambers, the time spent in each, the latency to first enter the light chamber, and the distance travelled in each chamber were recorded using a video camera attached to a computer and calculated by the Image LD program.

#### Open field test

Locomotor activity was measured using an open field test. Each mouse started the test in the corner of the open field apparatus (40 × 40 × 30 cm; Accuscan Instruments, O’hara & Co., LTD.). The chamber of the test was illuminated at 100 lux. Total distance (cm), total move time, and time spent in the center area were recorded.

#### Social interaction test

Sociability was measured using the open field apparatus. Mice of the same genotype were placed at diagonal corners in an open field apparatus in small baskets. After waiting for 1 minute, the behavior was recorded for 10 minutes after the mice exited from the baskets at the same time. The time to first contact, the total number of contacts, and the total time of sociable behavior were analyzed. All data was analyzed by a blind observer. The total sociable behavior time was measured excluding contact of less than one second. The test was performed with different pairs for a period of 2 days, and the average value for the 2 days was calculated. Mice were paired such that the weight difference was 2 g or less.

#### Elevated plus maze test

The elevated plus maze consisted of two open arms (25 × 5 cm) and two closed arms of the same size with 15 cm high transparent walls (O’hara & Co., LTD.). The behavior testing room (170 × 210 × 200 cm) was soundproof, and the illumination level was maintained at 100 lux. Each mouse was placed in the central square of the maze with its head pointed toward a closed arm. Data were recorded for 10 min using a video camera attached to a computer and were calculated with the Image EP program. The number of entries into each arm, total arm entries and the time spent in the open arms were recorded.

#### Rotarod test

Balance and motor ability were confirmed by the rotarod (BrainScience.idea.co., ltd). After placing the mouse in the rotating lane, the speed was changed from 3 to 30 rpm and the time until the mouse fell was measured. When one half of the body of the mouse was detached from the rods, it was measured as if it had fallen. The test was conducted for 3 consecutive days and 3 times a day.

#### Tail suspension test

Depression was investigated using a tail suspension test. The device used was the same as the fear test described next. The mouse was hung a hook with tape, and observed for 10 minutes. Recordings were made with a camera attached to the device and the freezing percentage was measured with software (O’hara & Co., LTD.). Freezing percentages were compared during the time elapsed excluding the first 3 minutes.

#### Fear condition test

On the 1st day, each mouse was placed into a test chamber (26 × 34 × 29 cm) with a stainless-steel grid floor inside a sound-attenuated chamber and allowed to explore freely for 120sec. A 60 db white noise acting as a conditioned stimulus (CS) was presented for 30 seconds, followed by a mild foot shock (2 seconds, 0.5 mA) acting as an unconditioned stimulus (US). Two more CS-US pairings were given at a stimulus interval of 2 minutes. On the 2nd day, a context test was performed in the same chamber as the conditioning test and data were recorded for 2 min. On the 3rd day, a cued test was performed in another testing chamber. The same 60 db white noise as on the 1st day was given for 30 s and data were recorded for 2 min. The entire test was recorded by a video camera attached to a computer. In each test, the movement freezing percentage and total move distance were calculated automatically by the Image FZ program.

### Immunohistochemistry

Mouse adult brains were fixed in 4% paraformaldehyde (PFA: Nacalai tesque), incubated in 10% and 25% sucrose at 4℃ overnight each and finally embedded in OCT compound (Sakura Finetek). The OCT blocks were sectioned at a 13-μm thickness with a cryostat (MICROTOME) and mounted on Superfrost Micro Slides (Matsunami Glass). The cryosections were fixed in 4% PFA for 10 min at room temperature and washed three times with PBS for 5 min. For antigen retrieval, the sections were boiled in 0.01 M Citrate Buffer pH 4.0 at 98℃ for 30 min (for RTL8A/8B), 0.01 M Citrate Buffer pH 4.0 at 90℃ for 15 min (for NF-H (RNF402), and then immersed (dehydrated) in cold methanol at −30℃ 30min. After being air dried, the sections were blocked with 10% goat serum, 1% bovine serum albumin (BSA: Sigma Aldrich), and 0.1% Triton-X 100 (WAKO) in PBS at room temperature for 1 hr.

For the staining, an anti-RTL8A/8B antibody (SCRUM 1:1000), and Anti Neurofilament antibody (NF-H (RNF402, Santa Cruz 1:200) were used as the primary antibodies and were prepared in 1% BSA and 0.1% Triton-X 100 in PBS at 4℃ overnight. The second antibody was Biotin-αRabbit-IgG (VECTOR STAIN 1:200). Nuclei were stained with Nuclear fast red. The slides were mounted with malinol (MUTO Pure Chemicals, Tokyo, Japan). The images were captured using a BIOREVO microscope (KEYENCE).

### Immunofluorescence staining using floating frozen section

Mouse brains were fixed and embedded in OCT blocks as described in the immunochemistry section. The OCT blocks were thinly sliced at 50 µm, floated in a glass bottom dish with 2 ml of PBS and washed for 10 min. PBS was removed and 2 ml of 0.1 M PBS/0.3% Triton was added and washed for 15 min three times. The sections were blocked with 10% donkey serum, 5% bovine serum albumin (BSA: Sigma Aldrich), and 0.1% Triton-X 100 (WAKO) in PBS at room temperature for 1 hr. For the staining, an anti-RTL8A/8B (SCRUM 1:2000) and anti-MAP2 (ab5392 1:20000) antibodies were used as the primary antibodies and were prepared in 5% BSA and 0.1% Triton-X 100 in PBS at 4℃ two overnight. Second antibodies used were the Alexa Fluor 488 donkey anti-Mouse IgG (H+L) (Life technologies, A21203,1:2000) and Alexa Fluor 488 donkey anti-Rabbit IgG (H+L) (Jackson ImmunoResearch, 711-545-152, 1:2000) and were prepared in 5% BSA and 0.1% Triton-X 100 in PBS at 4℃ overnight. Nuclei were stained with DAPI (VECTOR STAIN, 1:1000). The slides were mounted with VECTERSHIED Hardset antifade mounting medium (VECTOR STAIN). The images were captured using a confocal laser microscope TCS SP8 (Leica).

### Immunofluorescence staining using paraffin embedded samples

Mouse adult brains were fixed overnight using 4% paraformaldehyde (PFA: Nacalai tesque), and soaked in 70% ethanol at 4℃ overnight, then dehydrated in 70%, 80%, 90% ethanol for 1 hr each, 100% ethanol for 2 hr twice and 3hr once. They were embedded in paraffin wax after treatment with Gnox (Genostaff, GN04) for 2 hr twice, Gnox and paraffin for 2 hr, and paraffin for 3 hr and 4.5 hr. The paraffin blocks were sectioned at a 5μm thickness with a microtome and mounted on Superfrost Micro Slides (Matsunami Glass). For antigen retrieval, the sections were boiled in 0.01 M Citrate Buffer (pH 6.0) at 90℃ for 15 min then immersed (dehydrated) in cold methanol at 30℃ for 30min. After being air dried, the sections were treated with 10% goat serum, 1% bovine serum albumin (BSA: Sigma Aldrich), and 0.1% Triton-X 100 (WAKO) in PBS at room temperature for 1 hr.

For the immunofluorescence staining, anti-MAP2 (protein tech 17490-1-AP 1:200) antibody was reacted as the primary antibodies in 1% BSA and 0.1% Triton-X 100 in PBS at 4℃ overnight (approximately 16 hr). Second antibodies used were the Alexa Fluor 488 donkey anti-Rabbit IgG (H+L) (Jackson ImmunoResearch, 711-545-152, 1:1000). Nuclei were stained with Hoechst33342 (Sigma-Aldrich, 875756, 1:1000). The slides were mounted with VECTERSHIED Hardset antifade mounting medium (VECTOR STAIN). Neuronal areas were measured by a blind observer using ImageJ. The neuronal cells and nuclei were outlined manually with “Polygon selections” tool. The area of MAP2-positive neurons and the area of the nucleus were measured. Cytoplasmic area was calculated by subtracting nuclear area from cell area.

## Acknowledgments

The authors would like to thank Ms. Hitomi Takahashi (TMDU) for technical assistance in the embryo transfer experiment and Drs. Moe Kitazawa and Ayumi Matsuzawa (TMDU) for providing advice on the histological and bioinformatics experiments.

Pacific Edit reviewed the manuscript prior to submission.

## Author contributions

Y.F., M.I., H.S. T.E. Y.H. M.K. and F.I. performed the experiments and analyzed the data. M.I., R.O. H.K. and MK generated *Rtl8c*flox mice and R.O. and H. S generated *Rtl8a* and *Rtl8b* DKO mice. F.I. and T.K.-I. designed the study and Y.F., T.K.-I. and F.I. wrote the manuscript. All authors agree to be accountable for the content of the work.

---

**Fig. S1.**
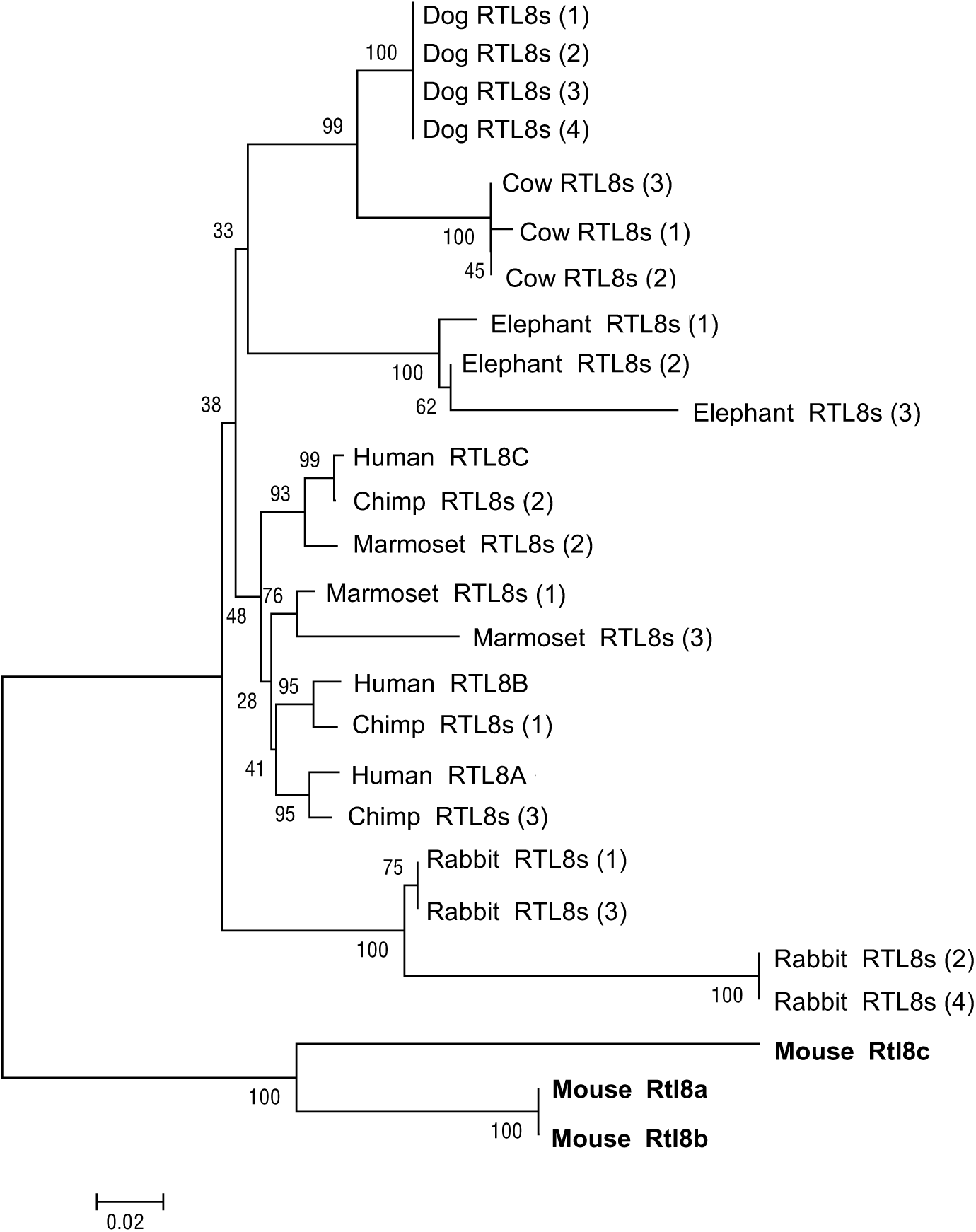
Evolutionary status of the *RTL8* genes in eutherians. In primates, human *RTL8A*, *8B* and *8C* are orthologous to Chimp *RTL8s* (1)-(3), but only marmoset *RTL8s* (2) is orthologous to human *RLT8C* and chimp *RTL8s* (2). In most cases, *RTL8* genes within each species exhibit a higher homology with each other than with those in other species, and thus are not in an orthologous relationship. The evolutionary history was inferred by using the Maximum Likelihood method based on the JTT

**Fig. S2.**
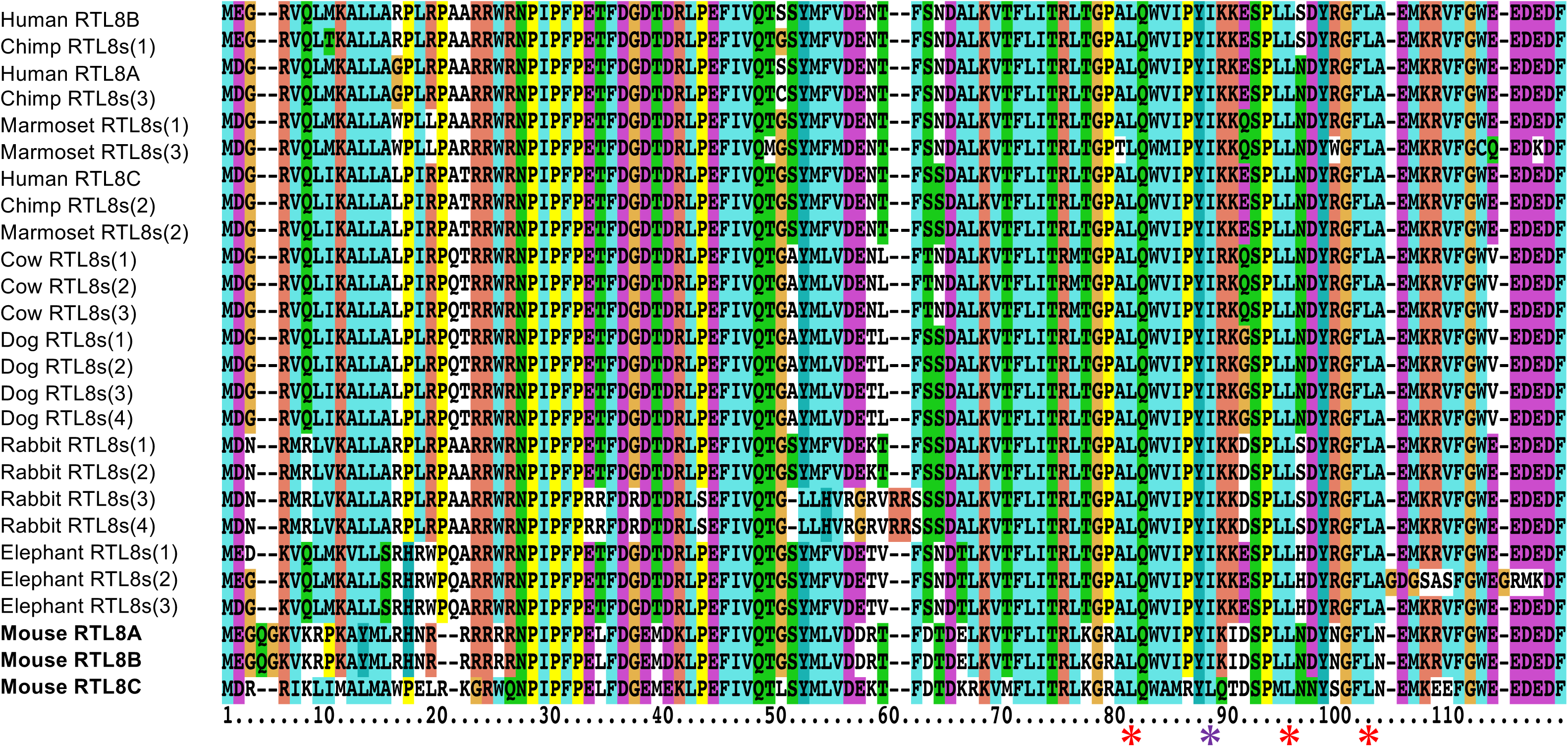
Comparison of amino acid sequences of RTL8A-C proteins in eutherians. A leucine zipper motif (3 red asterisks and 1 purple asterisk under the aa sequence is only present in mouse RTL8C (the bottom line) because the second leucine (purple) is replaced with isoleucine in the other RTL8A-C. Mouse RTL8A and 8B are unique because their N-terminal sequences contain the nuclear localization peptide-like sequences (see also Fig. 1B).

**Fig. S3.**
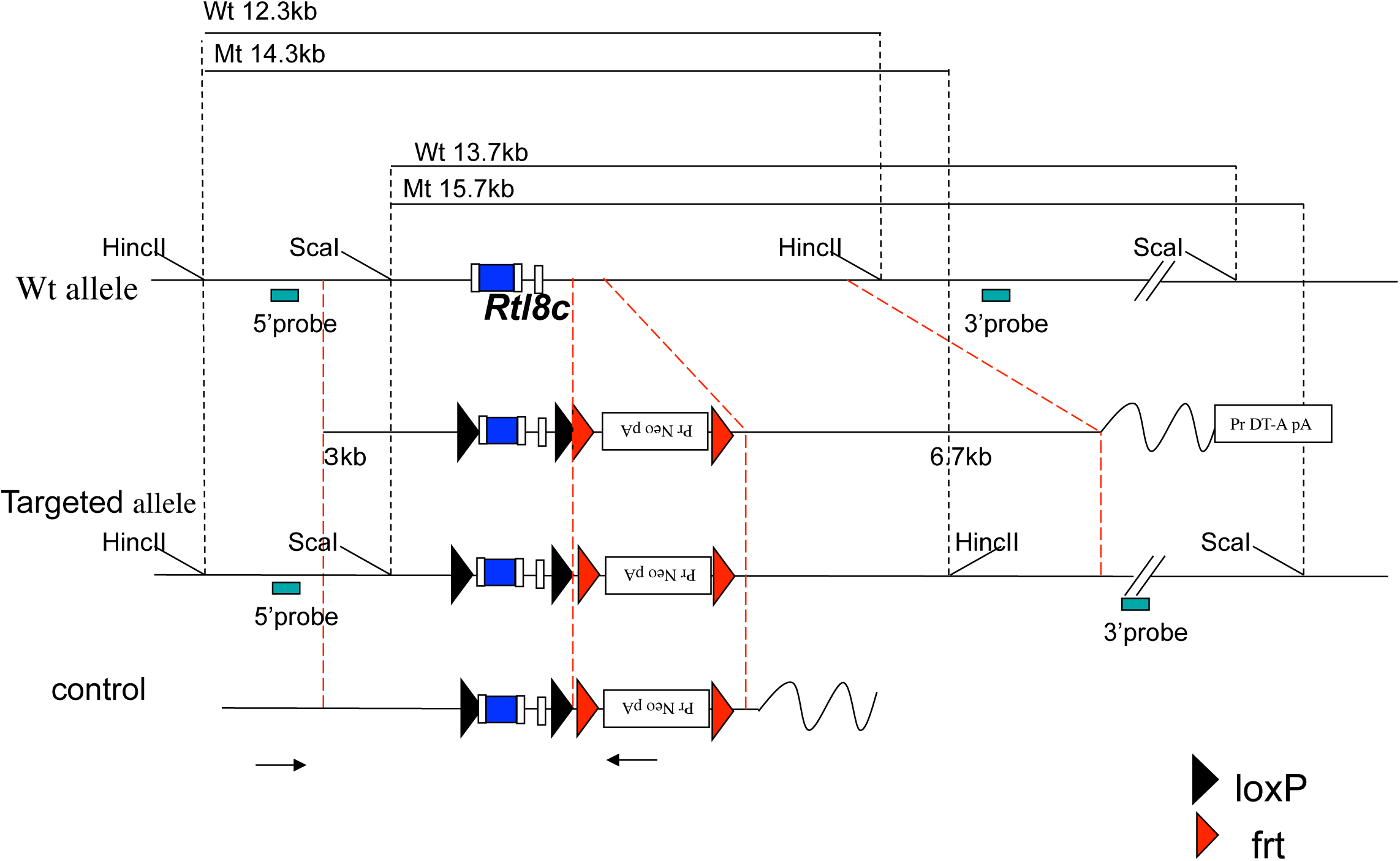
Construction of *Rtl8c* flox mouse. The *Rtl8c* flox targeting construct was generated using three genomic fragments, the 5′-arm (3.0 kb), middle arm containing *Rtl8c* exons and 3′-arm (6.5 kb). Two loxP sites were inserted upstream of exon 1 and downstream of exon 2, a frt-neo-frt cassette was inserted downstream of the latter loxP site. After linearizing, the construct was electroporated into TT2 ES cells. ES cells in which homologous recombination had occurred were injected into 8-cell stage embryos. Germ line transmission of the *Rtl8c* flox with the neomycin cassette (*Rtl8c* flox MT) was confirmed by Southern blot and PCR using the genome prepared from pups in which male *Rtl8c* flox chimeric mice had been crossed with female C57BL/6J. To remove the neo cassette, we mated the *Rtl8c* flox MT mice with CAG-FLP TG mice. The two white boxes represent *Rtl8c* exons and blue box in the exon 1 represent RTL8C ORF. The black and red arrowheads represent loxP and frt sites, respectively. The black arrows represent the PCR primers for screening of the targeted ES clones.

**Fig. S4.**
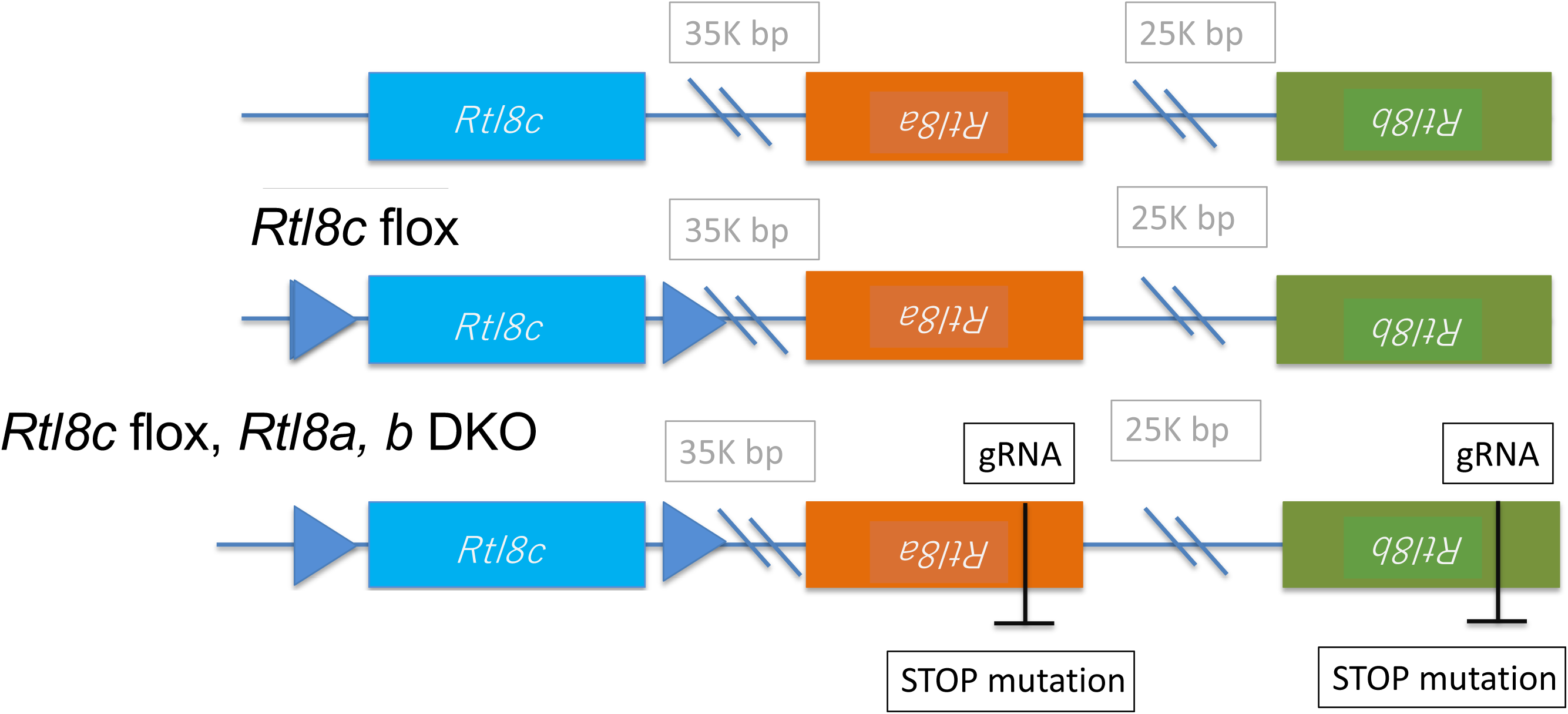
Strategy for making *Rtl8a* and *Rtl8b* DKO mouse. *Rtl8a* and *Rtl8b* are located tandemly while *Rtl8c* is reversed on the mouse X chromosome. First, *Rtl8c* flox mice were generated using TT2 ES cells (see Materials and Methods), then, *Rtl8a* and *Rtl8b* DKO mice were generated by integration of a stop codon in both *Rtl8a* and *Rtl8b*, as shown by the black bars, using CRIPR/Cas9.

**Fig. S5.**
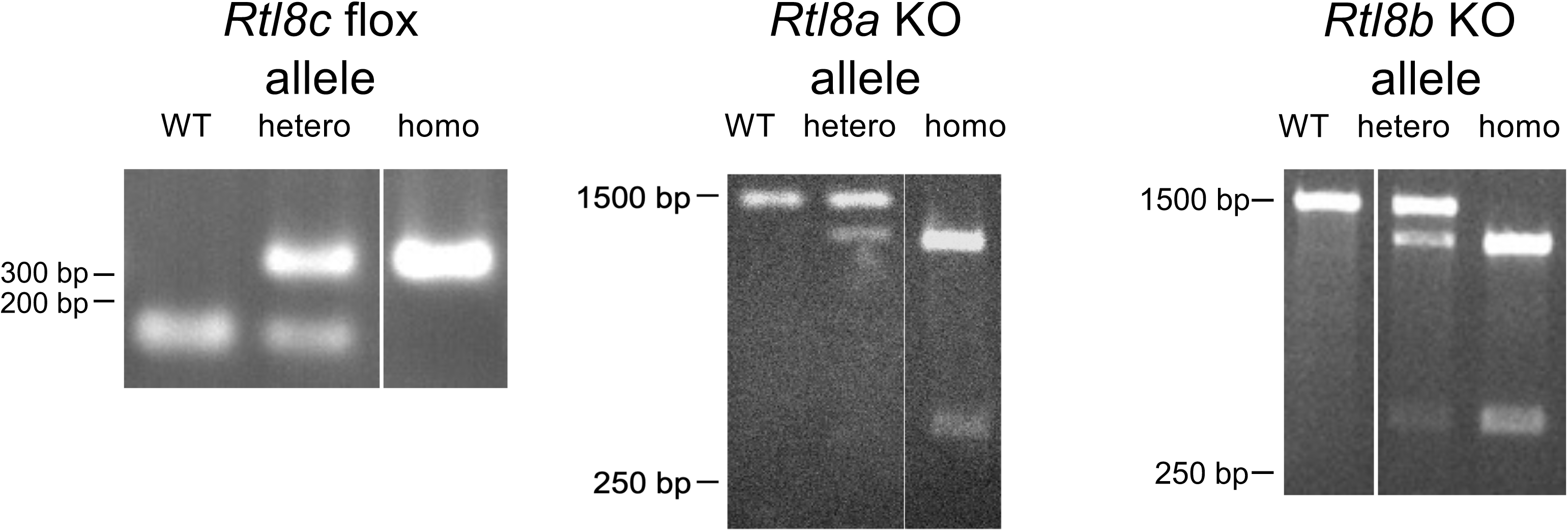
Genotyping the *Rtl8c* flox mice and the *Rtl8a* and *Rtl8b* DKO mice. Left: WT and *Rtl8c* flox alleles were detected as 169 and 349 bp bands in the PCR experiment, respectively. Middle: A *Rtl8a* KO allele with a stop mutation gave the same 1502 bp bands as WT, but were further digested by *Afi*II to give 1170 and 332 bp bands. Right: A *Rtl8b* KO allele with a stop mutation gave the same 1542 bp bands as WT, but were further digested by *Afi*II to give 1186 and 356 bp bands.

**Fig. S6.**
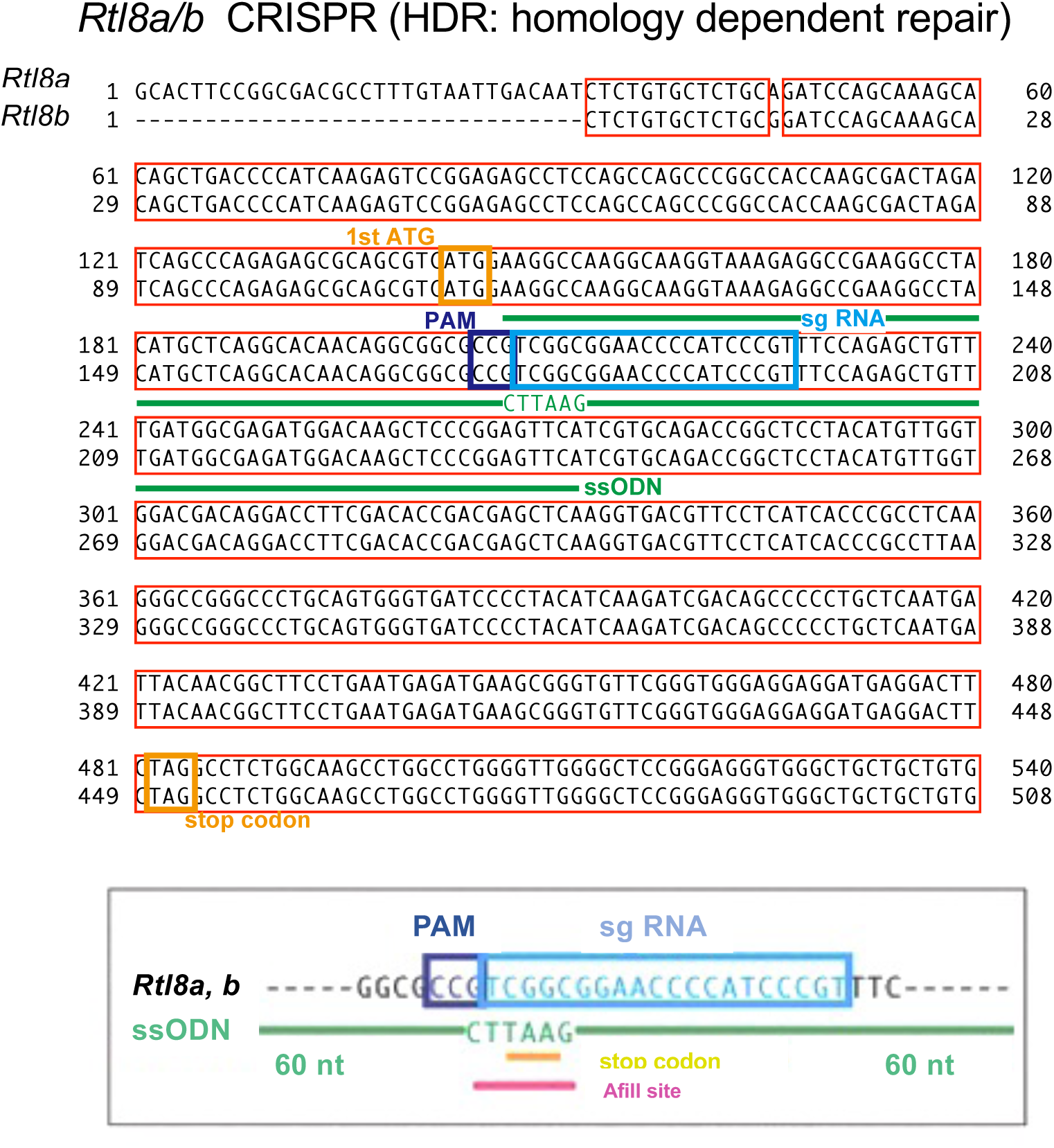
Construction of *Rtl8a* and *8b* DKO mouse. A stop codon was integrated in both *Rtl8a* and *Rtl8b* genes using the CRISPR/Cas9 method. The red and yellow boxes indicate the gene sequences of *Rtl8a* and *Rtl8b* as well as the start and stop codons, respectively. The light blue and purple box indicate the sgRNA and sPAM sequences, respectively. Pink line indicates the *Afl*II recognition site. ssODNs: single-stranded oligodeoxynucleotides.

**Fig. S7.**
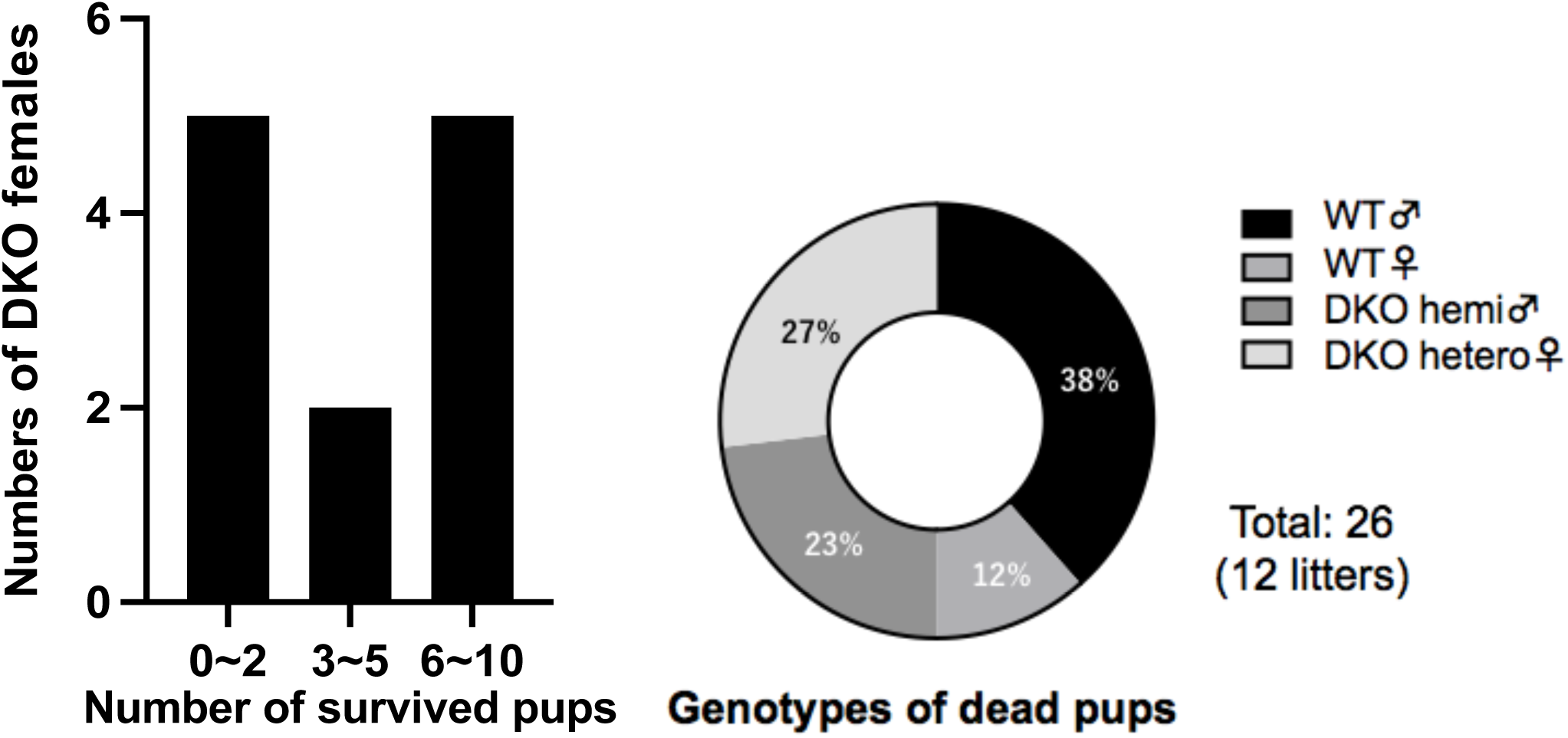
One half of the hetero DKO mothers failed to take proper care of their pups. Left: In most cases, hetero DKO mothers delivered 6-10 pups, but most pups were lost in half of the cases while most were viable in the other half. Right: Both male and female pups with WT and DKO genotypes were included in dead pups.

**Fig. S8.**
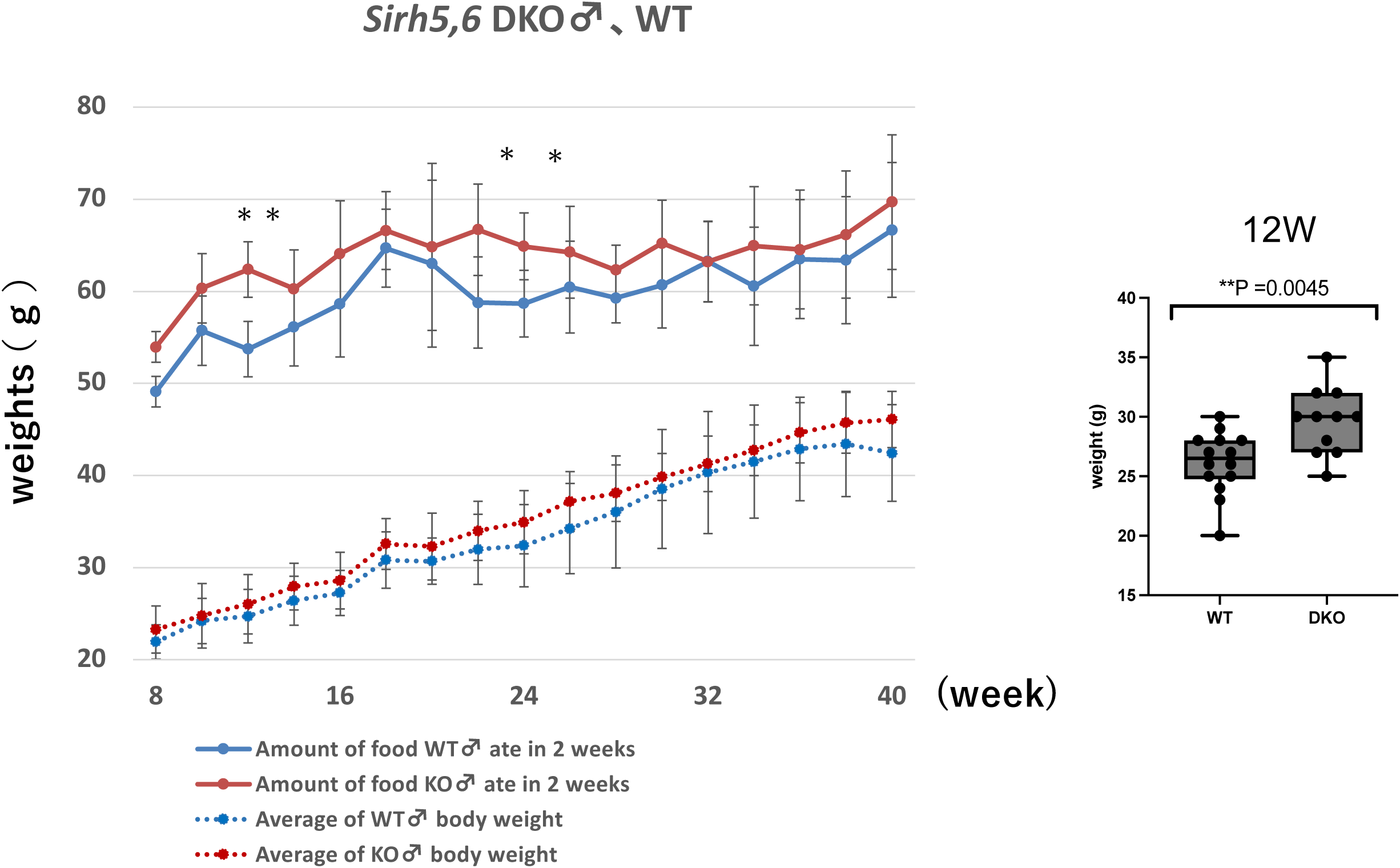
Growth curve of the WT and hemi DKO male mice. **(A)** The body weight (bottom) and food intake every two weeks (top) for the WT (n = 5) and hemi DKO male (n = 5) mice are shown. Blue: WT, Red: DKO. * P < 0.05. **(B)** The body weights of the WT (n = 14) and hemi DKO male (n = 11) mice at 12 weeks are shown. **P < 0.01.

**Fig. S9.**
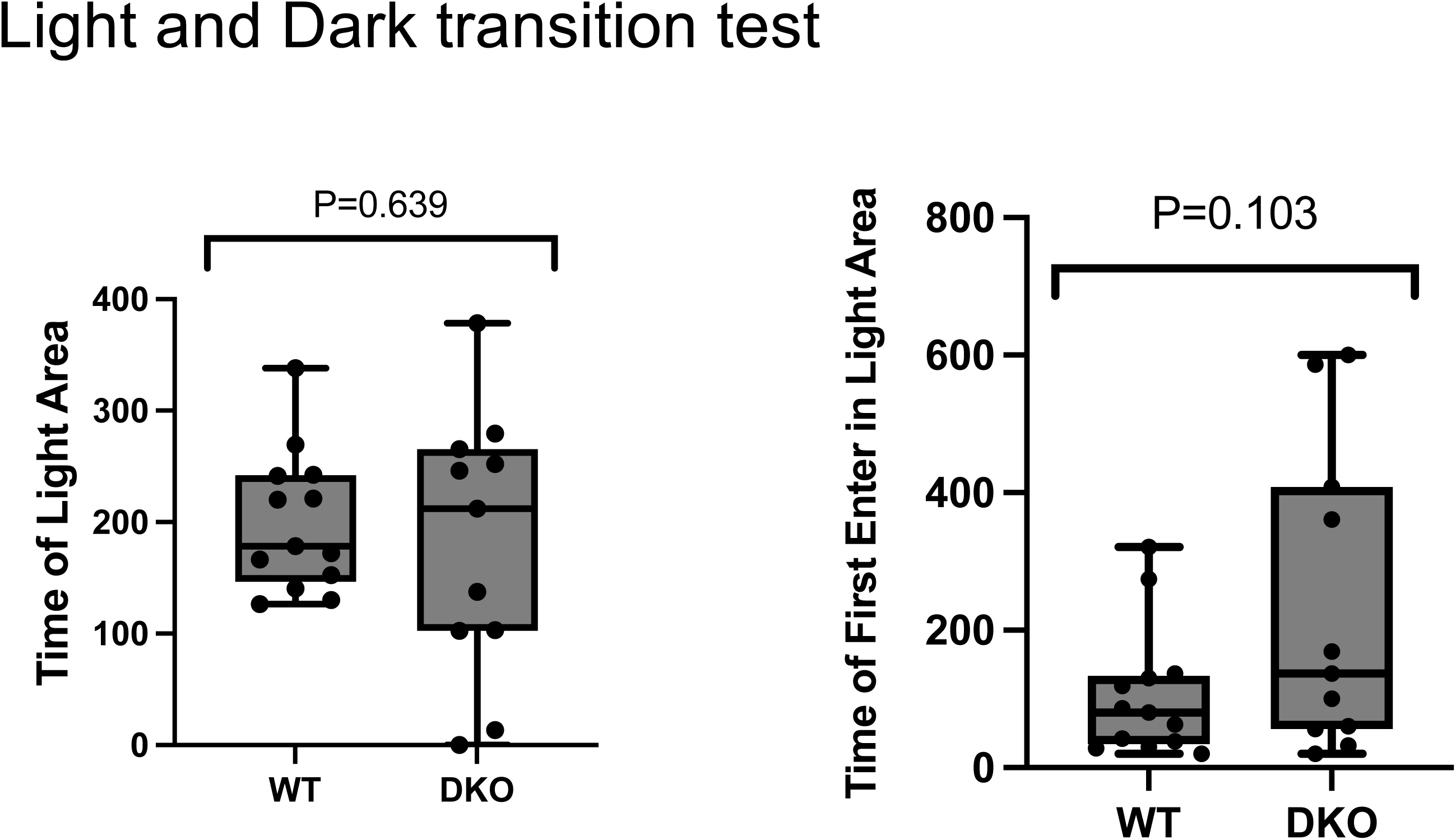
Light/Dark transition test. Results of Light/Dark transmission test except for those already shown in Fig. 2B. WT: n = 14, DKO: n = 11.

**Fig. S10.**
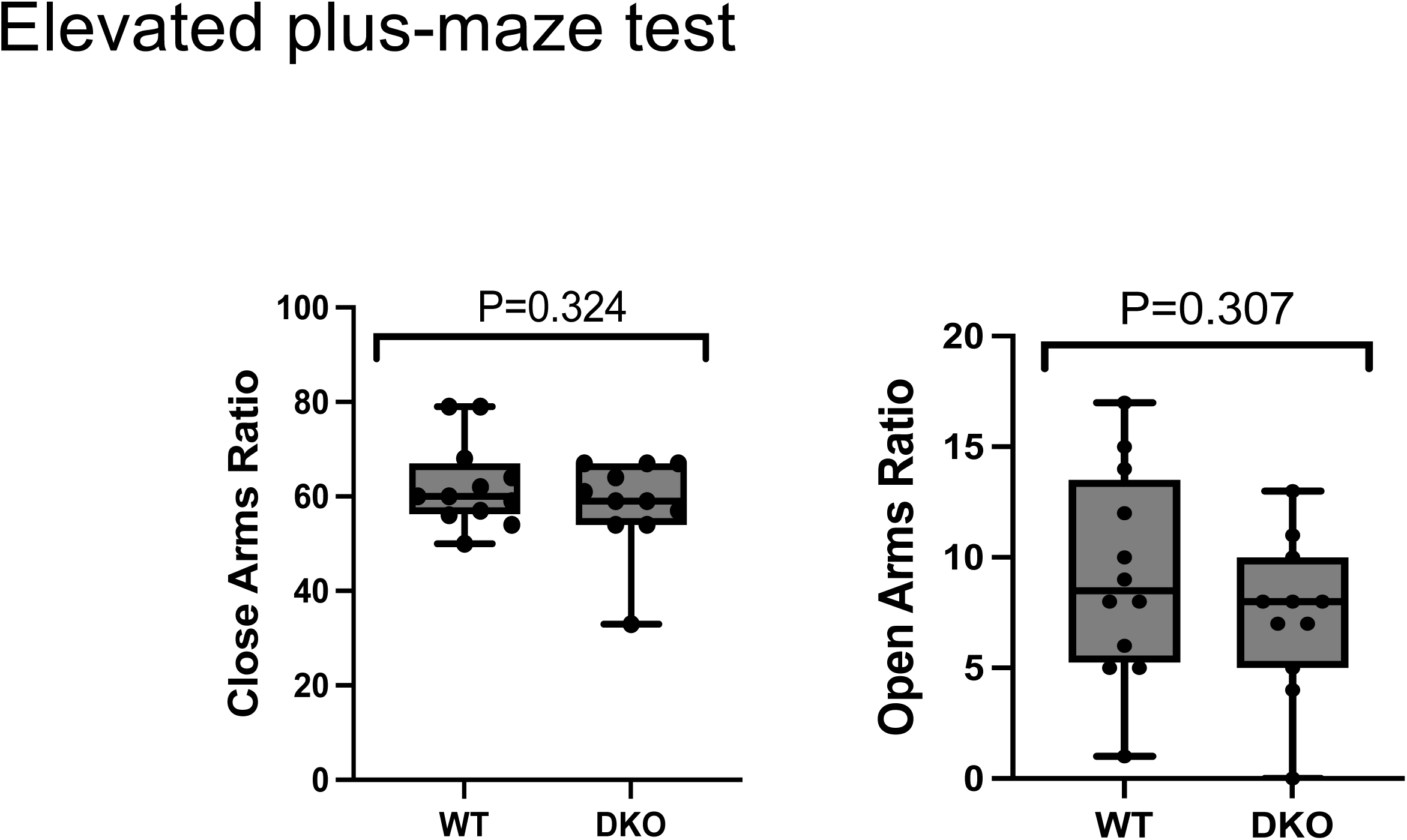
Elevated plus-maze test. Results of Elevated plus-maze test except for those already shown in Fig. 2B. WT: n = 12, DKO: n = 11.

**Fig. S11.**
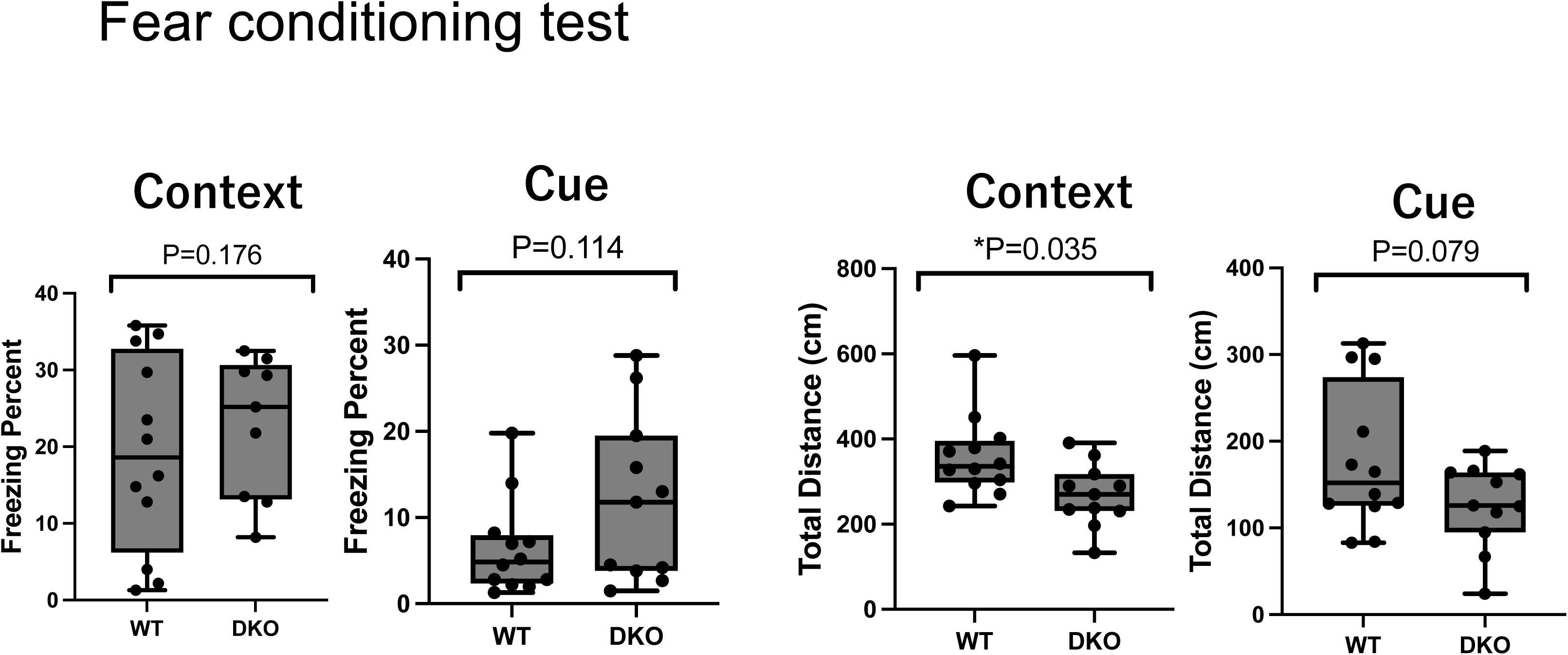
Fear conditioning test. Results of the Fear conditioning test are shown, WT: n = 12, DKO: n = 11.

**Fig. S12.**
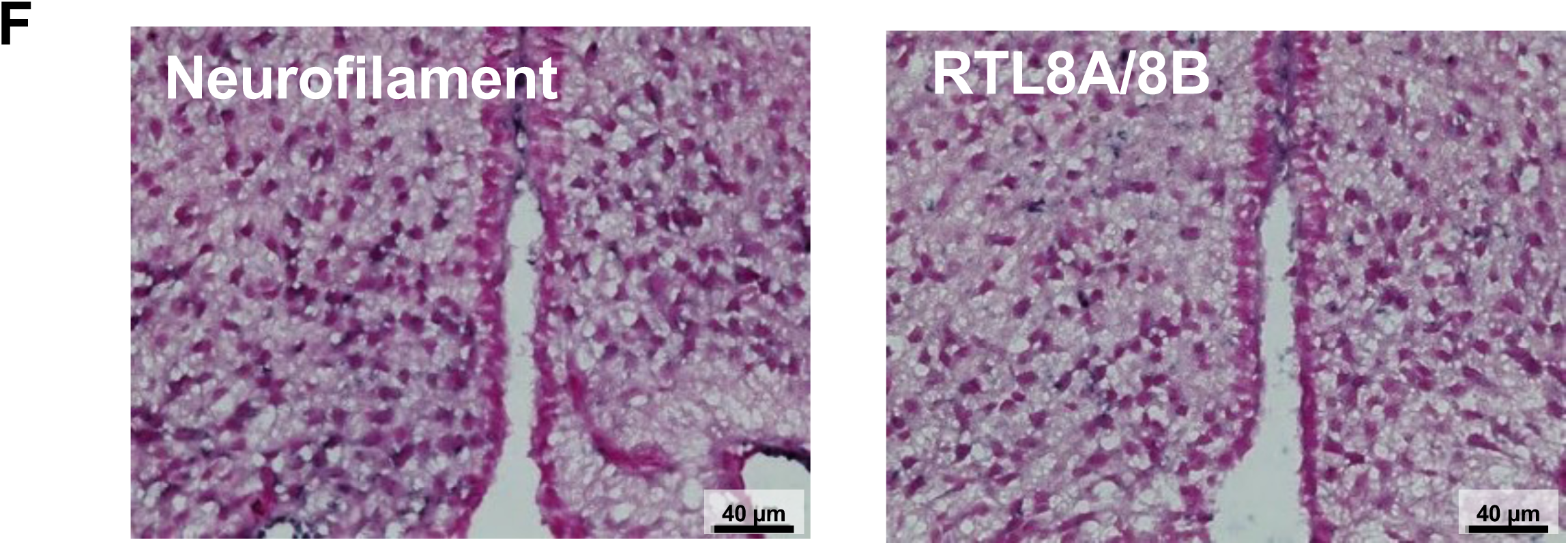
Immunostaining of mRTL8A/8B and Neurofilament. Consecutive two frozen sections of the WT mice (at 10 w) were used for the immunostaining of mRTL8A/8B and Neurofilament.

**Fig. S13.**
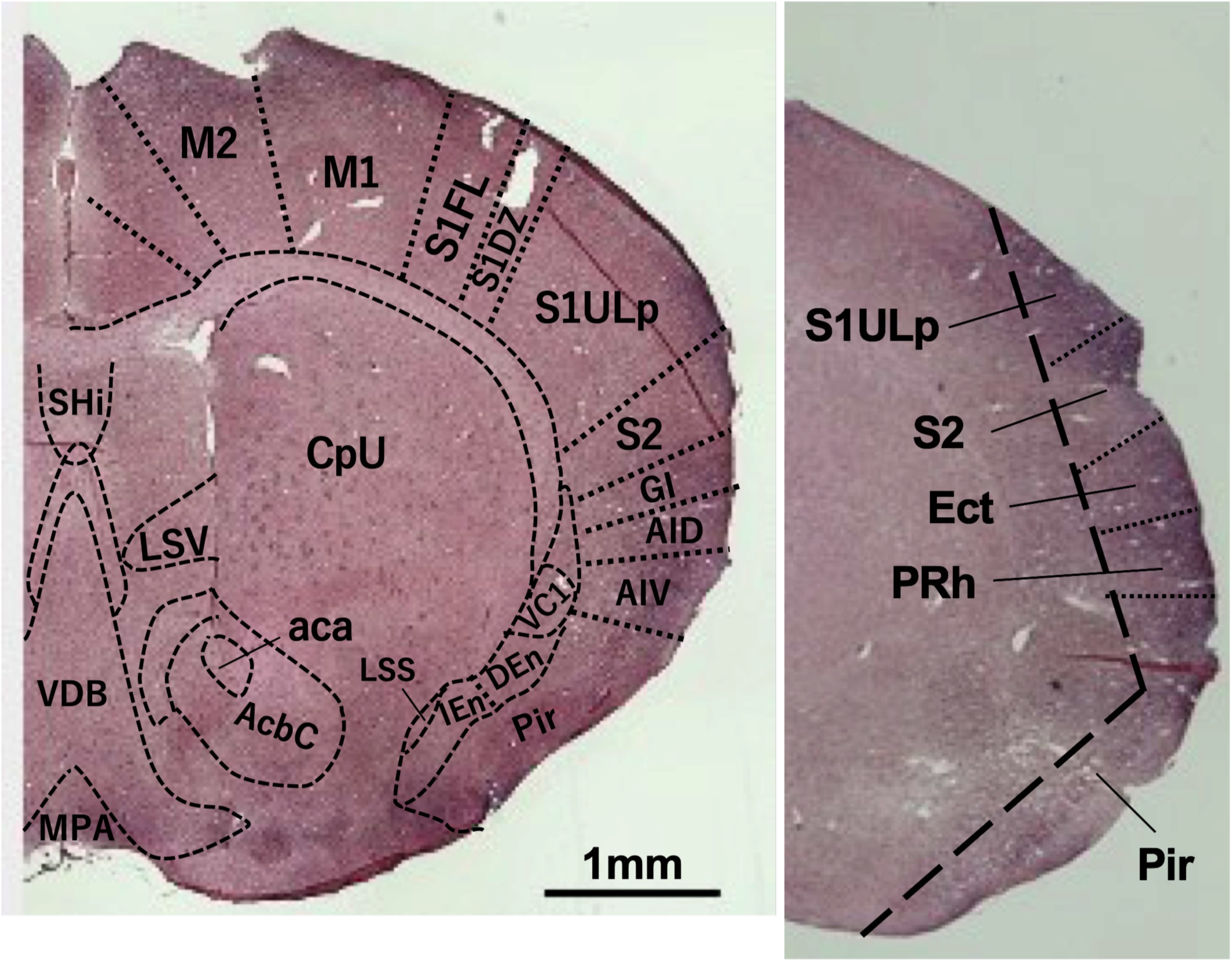
Immunostaining of RTL8A and RTL8B proteins in the striatum and temporal lobe. Coronal section images of WT mice showing staining in the striatum region (left: 10 w) and temporal lobe (right: 20 w). aca: anterior commissure, anterior part, AcbC: accumbens nucleus, core (ventral striatum), AID: agranular insular cortex, dorsal part, AIV: agranular insular cortex, ventral part, CpU: caudate putamen (dorsal striatum), DEn: dorsal endopiriform claustrum, Ect : ectorhinal cortex, GI: granular insular cortex, IEn: intermediate endopiriform claustrum, MPA: medial preootic area, M1: primary motor cortex, M2: secondary motor cortex, LSS: lateral stripe of the striatum (ventral stiatum), LSV: lateral septal nucleus, ventral part, Pir: piriform cortex, PRh : perirhinal cortex, S1DZ : primary somatosensory cortex, dysgranular zone, S1FL: primary somatosensory cortex, forelimb region, S1ULp: primary somatosensory cortex, upper lip region, S2: secondary somatosensory cortex, SHi: septohippocampal nucleus, VC1: ventral part of claustrum, VDB: nucleus of the vertical limb of the diagonal band.

**Fig. S14.**
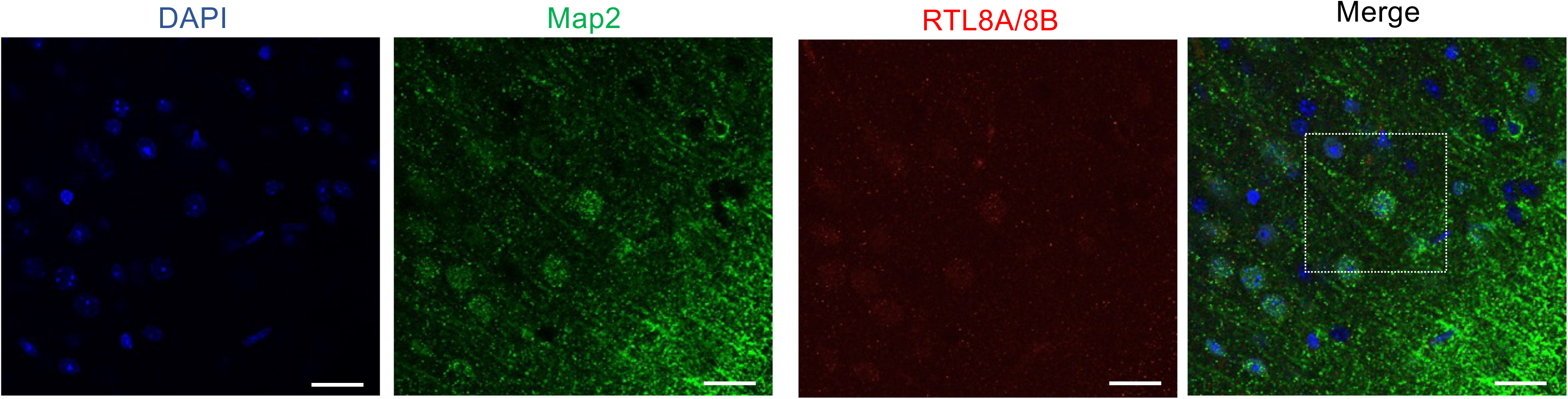
Co-immunofluorescence staining images of MAP2 and RTL8A/8B in the FrA region in WT. Lower magnification images of the Fig. 3F. A dashed square region shown in the right figure is enhanced in the Fig. 3F. Green: MAP2, Red: mRTL8A/8B blue: DAPI. Scale bars: 25 μm.

**Fig. S15.**
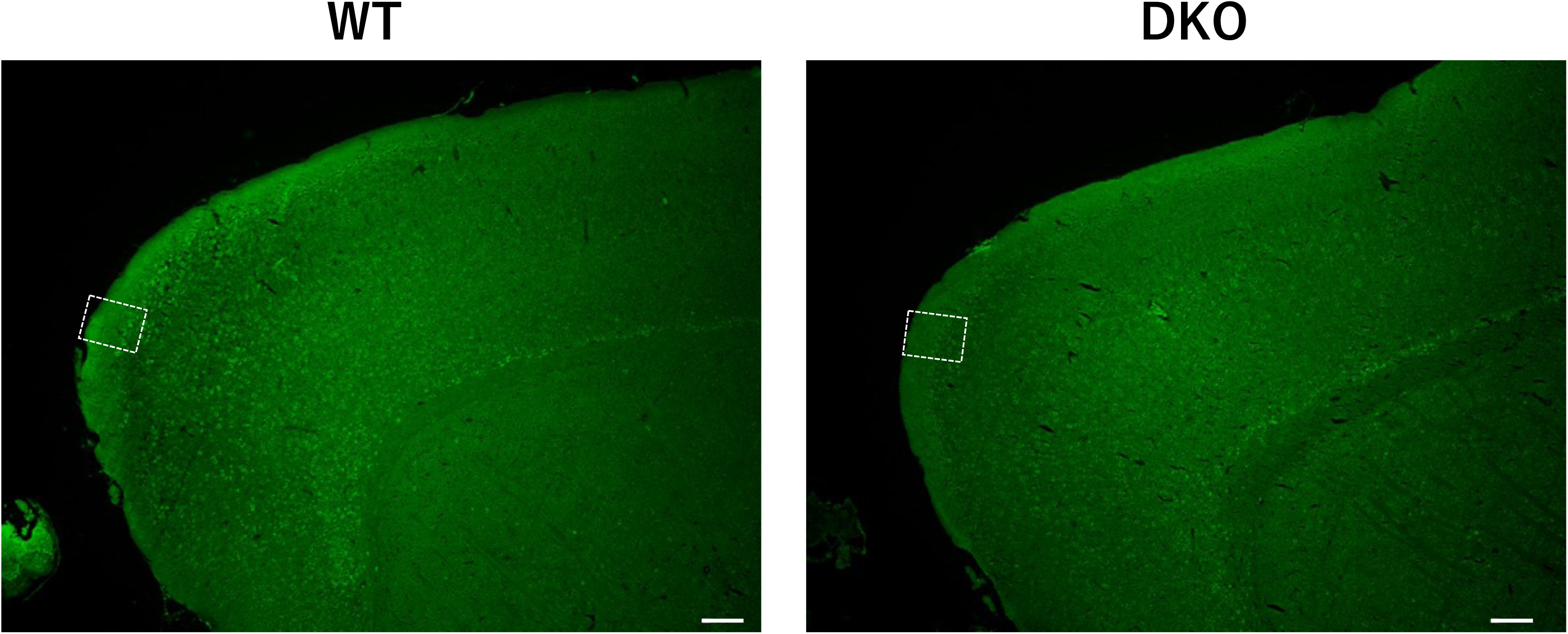
Fluorescence immunostaining images of the FrA region in WT and DKO mice. The areas surrounded by white squares are enlarged in Fig. 5A. Green: MAP2. Scale bar: 200 μm.

**Fig. S16.**
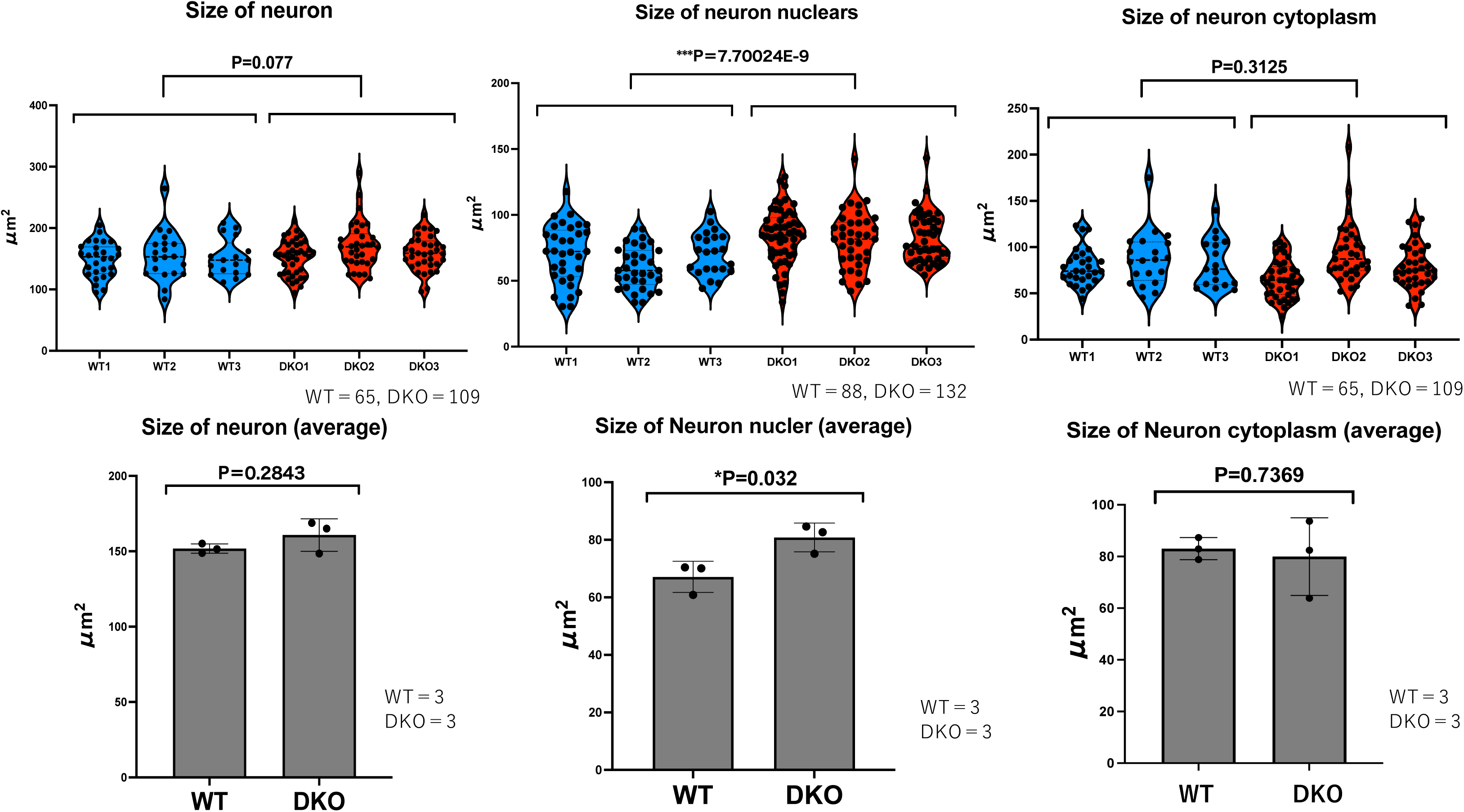
The sizes and average sizes of neurons, neuronal nuclei and neuronal cytoplasm in the FrA region. Top: Each of three individuals of the WT and DKO data in Fig. 5B were presented. Bottom: Their averages were presented statistically analyzed with Student’s t-test. *P < 0.05, **P < 0.01, ***P< 0.001.

**Fig. S17.**
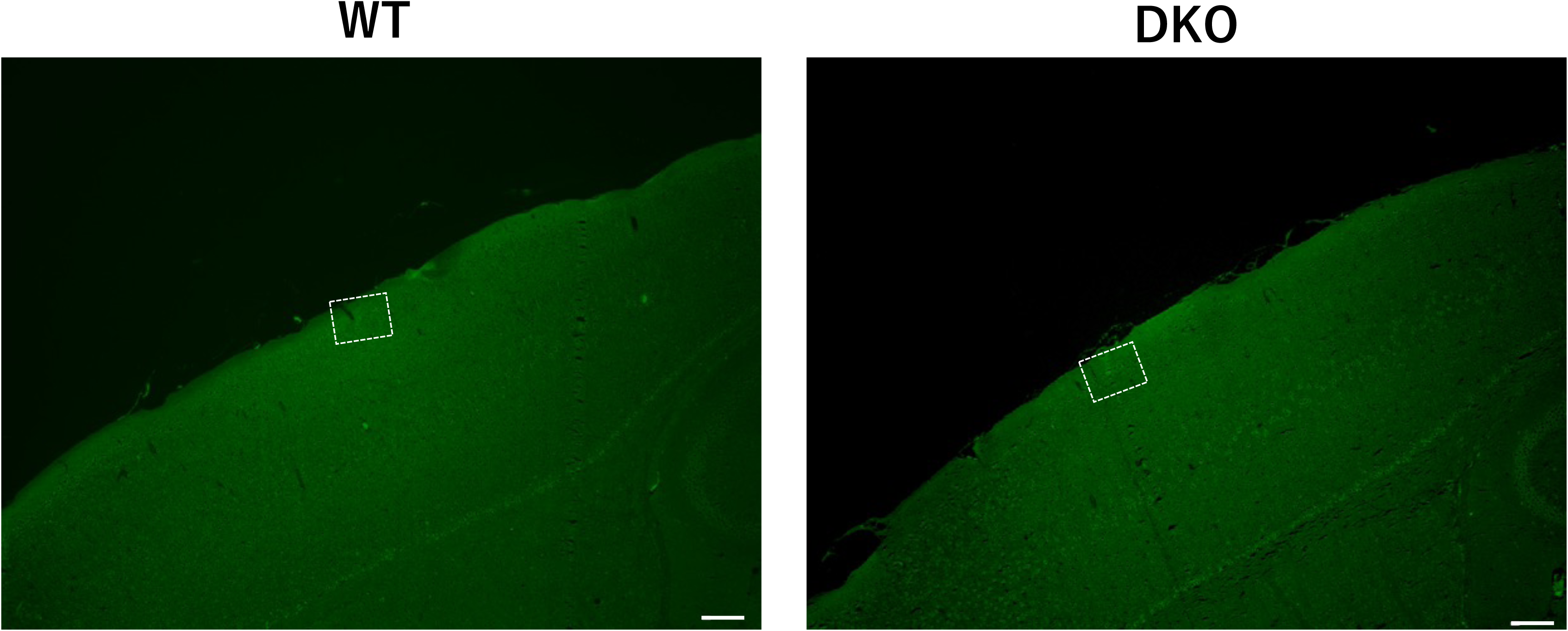
Fluorescence immunostaining images of the S1Sh region in WT and DKO mice. The areas surrounded by white squares are enlarged in Fig. 5D. Green: MAP2. Scale bar: 200 μm.

**Fig. S18.**
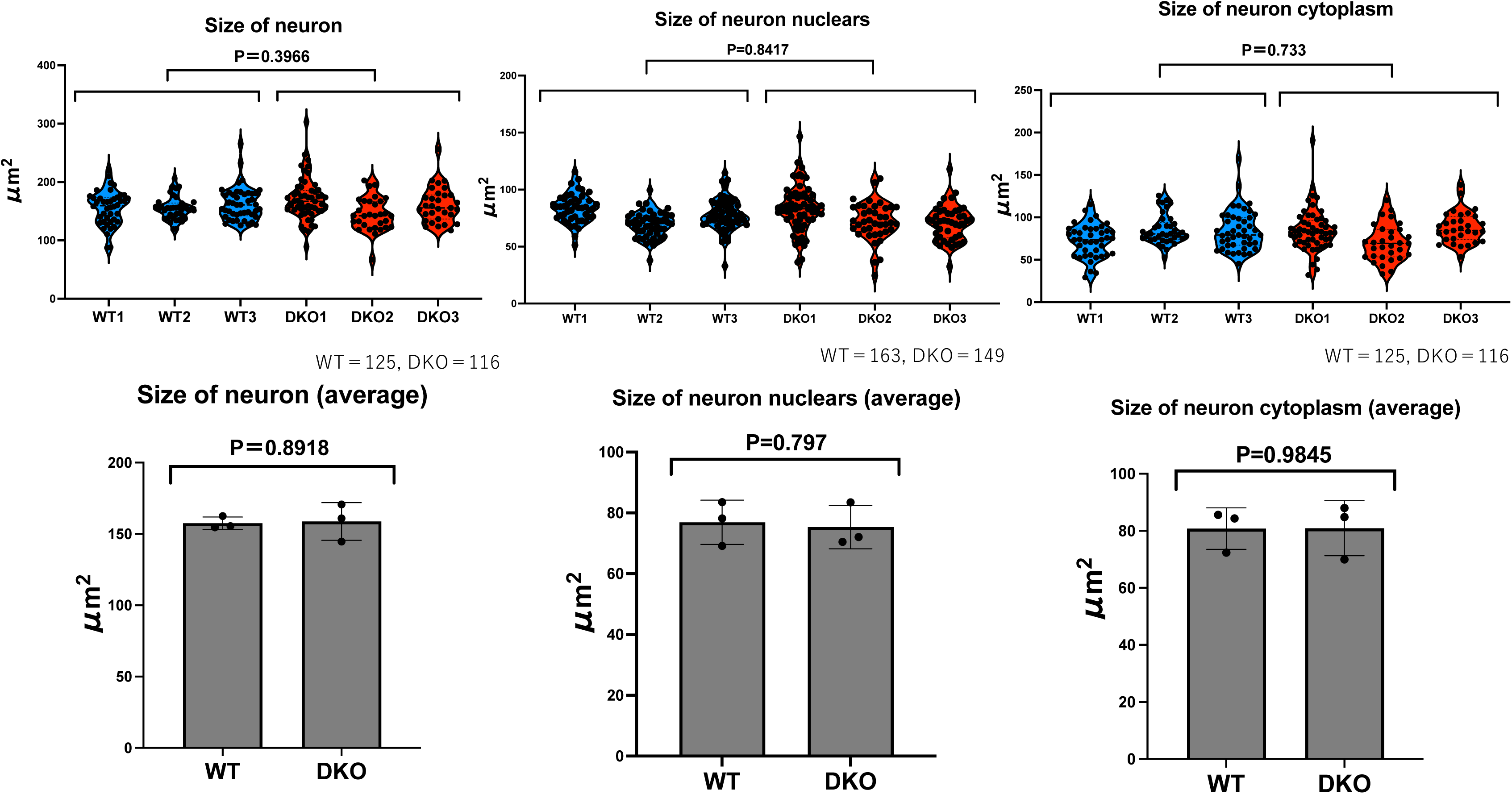
The sizes and average sizes of the neurons, neuronal nuclei and neuronal cytoplasm in the S1Sh region. Top: Each of three individuals of the WT and DKO data in Fig. 5D were presented. Bottom: Their averages were presented statistically analyzed with Student’s t-test. *P < 0.05, **P < 0.01, ***P< 0.001.

## References

Atasoy, D. et al. (2012) Deconstruction of a neural circuit for hunger. Nature 488, 172– 177.

Brandt J, Schrauth S, Veith AM, Froschauer A, Haneke T, Schultheis C, Gessler M, Leimeister C, Volff JN (2005) Transposable elements as a source of genetic innovation: expression and evolution of a family of retrotransposon derived neogenes in mammals. Gene 345: 101–111

Broschard, M. B. et al. (2021) Prelimbic cortex maintains attention to category-relevant information and flexibly updates category representations. Neurobiol Learn Mem 185, 107524.

Charlier C, Segers K, Wagenaar D, Karim L, Berghmans S, Jaillon O, Shay T, Weissenbach J, Cockett N, Gyapay G, Georges M (2001) Human–ovine comparative sequencing of a 250-kb imprinted domain encompassing the callipyge (clpg) locus and identification of six imprinted transcripts: DLK1, DAT, GTL2, PEG11, antiPEG11, and MEG8. Genome Res 11: 850–862

Chou, M. Y. et al. (2022) RTL1/PEG11 imprinted in human and mouse brain mediates anxiety-like and social behaviors and regulates neuronal excitability in the locus coeruleus. Hum Mol Genet 31, 3161–3180.

Hare, T. A. et al. (2009) Self-control in decision-making involves modulation of the vmPFC valuation system. Science 324, 646– 648.

Hare, B. D. and Duman, R. S. (2020) Prefrontal cortex circuits in depression and anxiety: contribution of discrete neuronal populations and target regions. Mol Psychiatry 25, 2742–2758.

Imakawa, K., et al. (2022) Endogenous Retroviruses and Placental Evolution, Development, and Diversity. Cells 11, 2458.

Ioannides, Y. et al. (2014) Temple syndrome: Improving the recognition of an underdiagnosed chromosome 14 imprinting disorder: An analysis of 51 published cases. J Med Genet 51. 495–501.

Irie, M. et al. (2015) Cognitive Function Related to the Sirh11/Zcchc16 Gene Acquired from an LTR Retrotransposon in Eutherians. PLoS Genet 11, e1005521.

Irie, M., et al. (2022) Retrovirus-derived *RTL5* and *RTL6* genes are novel constituents of the innate immune system in the eutherian brain. Development 149, dev200976.

Kagami, M., et al. (2008) Deletions and epimutations affecting the human 14q32.2 imprinted region in individuals with paternal and maternal upd(14)-like phenotypes. Nat Genet 40, 237–242.

Kagami, M., et al. (2015) Comprehensive clinical studies in 34 patients with molecularly defined UPD(14)pat and related conditions (Kagami-Ogata syndrome). Eur J Hum Genet 23, 1488–1498.

Kaneko-Ishino, T. and Ishino, F. (2012) The role of genes domesticated from LTR retrotransposons and retroviruses in mammals. Front Microbiol 3, 262.

Kaneko-Ishino, T. and Ishino, F. (2015) Mammalian-specific genomic functions: Newly acquired traits generated by genomic imprinting and LTR retrotransposon-derived genes in mammals. Proc Jpn Acad Ser B Phys Biol Sci 91, 511–538.

Kitazawa, M. et al. (2020) Deficiency and overexpression of *Rtl1* in the mouse cause distinct muscle abnormalities related to the Temple and Kagami-Ogata syndromes. Development 147, dev185918.

Kitazawa, M., et al. (2021) The role of eutherian specific *RTL1* in the nervous system and its implications for the Kagami-Ogata and Temple syndromes. Genes Cells 26, 165–179.

Lanfray, D. and Richard, D. (2017) Emerging Signaling Pathway in Arcuate Feeding Related Neurons: Role of the Acbd7. Front Neurosci 11, 328.

Lim, E. T. et al. (2013) Rare complete knockouts in humans: population distribution and significant role in autism spectrum disorders. Neuron 77. 235–242.

Merkestein, M., et al. (2014) GHS-R1a signaling in the DMH and VMH contributes to food anticipatory activity. Int J Obes 38, 610–618.

Mohan, H. M., et al. (2022) RTL8 promotes nuclear localization of UBQLN2 to subnuclear compartments associated with protein quality control. Cell Mol Life Sci 79,176.

Nakayama, D., et al. (2015) Frontal association cortex is engaged in stimulus integration during associative learning. Curr Biol 25, 117–123.

Naruse, M., et al. (2014) *Sirh7/Ldoc1* knockout mice exhibit placental P4 overproduction and delayed parturition. Development 141, 4763–4771.

Nicholls, R. D. and Knepper, J. L. (2001) Genome organization, function, and imprinting in Prader-Willi and Angelman syndromes. Annu Rev Genomics Hum Genet 2, 153–175.

Ono, R., et al. (2001) A retrotransposon-derived gene, PEG10, is a novel imprinted gene located on human chromosome 7q21. Genomics 73, 232–237.

Ono, R. et al. (2006) Deletion of *Peg10*, an imprinted gene acquired from a retrotransposon, causes early embryonic lethality. Nat Genet 38, 101–106.

Ono, R., et al. (2015) Double strand break repair by capture of retrotransposon sequences and reverse-transcribed spliced mRNA sequences in mouse zygotes. Sci Rep 5, 12281.

Pandya, N. J., et al. (2021) Secreted retrovirus-like GAG-domain-containing protein PEG10 is regulated by UBE3A and is involved in Angelman syndrome pathophysiology. Cell Rep Med 2, 100360.

Pizzagalli, D. A. and Roberts, A. C. (2022). Prefrontal cortex and depression. Neuropsychopharmacology 47, 225–246.

Seabrook, L.T. and Borgland, S. L. (2020) The orbitofrontal cortex, food intake and obesity. J Psychiatry Neurosci 45, 304–312.

Sekita, Y. et al. (2008) Role of retrotransposon-derived imprinted gene, Rtl1, in the feto-maternal interface of mouse placenta. Nat Genet 40, 243–248.

Shrestha, P., et al. (2015) Layer 2/3 pyramidal cells in the medial prefrontal cortex moderate stress induced depressive behaviors. eLife 4, e08752.

Suyama, M., et al. (2006) PAL2NAL: robust conversion of protein sequence alignments into the corresponding codon alignments. Nucleic Acids Res. 34, W609–612.

Szczepanski, S. M. and Knight, R. T. (2014) Insights into human behavior from lesions to the prefrontal cortex. Neuron 83, 1002–1018.

Tamura, K., et al. (2011) MEGA5: Molecular Evolutionary Genetics Analysis Using Maximum Likelihood, Evolutionary Distance, and Maximum Parsimony Methods. Mol Biol Evol 28, 2751–2739.

Wang, H., et al. (2013) One-step generation of mice carrying mutations in multiple genes by CRISPR/Cas-mediated genome engineering. Cell 153, 910–918.

Whiteley, A. M. et al. (2021) Global proteomics of Ubqln2-based murine models of ALS. J Biol Chem 296, 100153.

Xu, B. and Yang, Z. (2013) PAMLX: a graphical user interface for PAML. Mol Biol Evol 30, 2723–2724.

Yagi, T., et al. (1993) A novel ES cell line, TT2, with high germline-differentiating potency. Anal Biochem 214, 70–76.

Youngson, N. A., et al. (2005) A small family of sushi-class retrotransposon-derived genes in mammals and their relation to genomic imprinting. J Mol Evol 61, 481–490.

Yousefvand, S. and Hamidi, F. (2020) Role of Paraventricular Nucleus in Regulation of Feeding Behaviour and the Design of Intranuclear Neuronal Pathway Communications. Int J Pept Res Ther 26, 1231–1242.

